# Reduced Levels of Lagging Strand Polymerases Shape Stem Cell Chromatin

**DOI:** 10.1101/2024.04.26.591383

**Authors:** Jonathan Snedeker, Brendon E. M. Davis, Rajesh Ranjan, Matthew Wooten, Joshua Blundon, Xin Chen

**Affiliations:** Department of Biology, The Johns Hopkins University, Baltimore, MD 21218, USA; Howard Hughes Medical Institute, Department of Biology, The Johns Hopkins University, 3400 North Charles Street, Baltimore, Baltimore, MD 21218, USA; Fred Hutchinson Cancer Research Center, Seattle, WA 98109-1024, USA

**Keywords:** germline stem cell, cellular differentiation, asymmetric cell division, DNA replication, chromatin fiber, DNA fiber, lagging strand synthesis, replication-dependent histone incorporation

## Abstract

Stem cells display asymmetric histone inheritance while non-stem progenitor cells exhibit symmetric patterns in the *Drosophila* male germline lineage. Here, we report that components involved in lagging strand synthesis, such as DNA polymerase α and δ (Polα and Polδ), have significantly reduced levels in stem cells compared to progenitor cells. Compromising Polα genetically induces the replication-coupled histone incorporation pattern in progenitor cells to be indistinguishable from that in stem cells, which can be recapitulated using a Polα inhibitor in a concentration-dependent manner. Furthermore, stem cell-derived chromatin fibers display a higher degree of old histone recycling by the leading strand compared to progenitor cell-derived chromatin fibers. However, upon reducing Polα levels in progenitor cells, the chromatin fibers now display asymmetric old histone recycling just like GSC-derived fibers. The old *versus* new histone asymmetry is comparable between stem cells and progenitor cells at both S-phase and M-phase. Together, these results indicate that developmentally programmed expression of key DNA replication components is important to shape stem cell chromatin. Furthermore, manipulating one crucial DNA replication component can induce replication-coupled histone dynamics in non-stem cells in a manner similar to that in stem cells.

**One Sentence Summary:** Delayed lagging strand synthesis regulates asymmetric histone incorporation.

## Introduction

Regarding metazoan development, an outstanding question is how cells take on distinct fates and have diverse functions even though they derive from one zygote. Cell fate is determined by selectively expressing a subset of the genome at the proper time, in the right place, and at the precise level. The unique gene expression program for each cell type is typically regulated by the epigenetic mechanisms, which refer to chromatin changes without alteration of the DNA sequences (*1–3*). Epigenetic mechanisms comprise DNA methylation, histone modifications, histone variants, as well as non-coding RNAs, among others. However, except for DNA methylation, how the epigenetic information is transferred through the active cell cycle in multicellular organisms remains largely unclear (*4*). Notably, these mechanisms could not only be responsible for maintaining epigenetic memory but also allow for epigenetic changes to diversify cell fates, which are essential for development, homeostasis and regeneration (*5–7*). One paradigmatic model to study cell fate decision is asymmetric cell division (ACD), through which one mother cell gives rise to two distinct daughter cells. Upon ACD, the genetic codes inherited by the two daughter cells are identical, whereas their epigenetic information can vary, allowing them to appear and function differently [reviewed by (*8–12*)]. Recently, it has been revealed that ACD can be induced by DNA damage in otherwise symmetrically dividing human cells, suggesting that changes on DNA strands *per se* could guide the cell division mode and potentially regulate the epigenetic inheritance pattern (*13*).

To investigate the histone inheritance pattern in ACD, a tag-switch strategy to differentially label preexisting (old) *versus* newly synthesized (new) histones has been developed and used to study the *Drosophila* adult stem cell systems. These studies reveal that old histones are selectively retained in the self-renewing stem cell, whereas new histones are enriched in the differentiating daughter cell during ACDs of male germline stem cells (GSCs) (*14, 15*) and intestinal stem cells (*16*). Notably, in the male germline lineage, old and new histones are inherited symmetrically during the symmetric divisions of the progenitor spermatogonial cells (SGs). Asymmetric histone inheritance has been proposed to involve a process with at least three steps: First, old and new histones are asymmetrically incorporated on the replicative sister chromatids, attributed by both strand-specific incorporation and biased replication fork movement, including increased unidirectional and asymmetric bidirectional fork progression in early-stage germ cells (*15, 17*). Then, the epigenetically distinct sister chromatids are differentially recognized and segregated during mitosis (*18*), leading to distinct “read-outs” in the resulting two daughter cells, such as their asynchronous S-phase initiation (*19*) and distinct interchromosomal interactions at a key “stemness” gene (*20*).

Despite this knowledge, two crucial questions still remain: First, what are the precise molecular mechanisms that ensure asymmetric histone incorporation at the individual replication forks? A series of studies have extensively explored the roles of DNA replication components in establishing the epigenomes in unicellular organisms, such as yeast (*21–25*), and symmetrically dividing cells, such as cultured mouse embryonic stem cells (*26–30*) and human cell lines (*31, 32*). These studies focus on how epigenetic information can be equally partitioned between sister chromatids and inherited symmetrically by the daughter cells [reviewed by (*4, 33–36*)]. Nevertheless, little is known about this process in asymmetrically dividing cells in multicellular organisms. Studies in mouse development demonstrate that asymmetric inheritance of H3R26me2 (*37*) or maternal chromosome-bound H3.3 and H3K9me2 (*38*) are essential for early embryogenesis, in contrast to the negative effects of asymmetric histone inheritance in yeast (*39–41*) and mouse embryonic stem cells (*28, 29*), emphasizing the importance to study this phenomenon in an organism-and context-dependent manner. Second, how are these mechanisms regulated in a stage-specific manner within the same adult stem cell lineage, such that histone inheritance is asymmetric in stem cells (e.g., GSCs) but symmetric in progenitor cells (e.g., SGs)? Here, we used the *Drosophila* male germline as a model system to address these questions.

## Results

### Differential expression of the lagging strand-enriched replication components in GSCs

To identify which factors could be responsible for stem cell-specific asymmetric histone inheritance, we performed a candidate gene screen using a series of CRISPR/Cas9-mediated knock-in lines with the <u>h</u>em<u>a</u>gglutinin (HA) tag at individual genes that encode distinct key replication machinery components. Intriguingly, the levels of proteins involved in lagging strand synthesis, such as DNA polymerase α and δ (Polα and Polδ), differ significantly between GSCs and SGs, with substantially reduced levels in GSCs compared to SGs (Fig. 1a). In contrast, a key component for leading strand synthesis, DNA polymerase ε (Polε), exhibits comparable levels between GSCs and SGs (Fig. 1a). On the other hand, the single-stranded DNA (ssDNA) binding protein Replication Protein-A 70 (RPA70), the largest subunit of the ssDNA-binding heterotrimeric complex (*42, 43*), displays the opposite trend with higher RPA levels in GSCs compared to SGs (Fig. 1a), using the RPA70-EGFP fusion protein expressed under the endogenous regulatory elements of the *rpa70* gene (*15, 44*).

**Figure 1:**
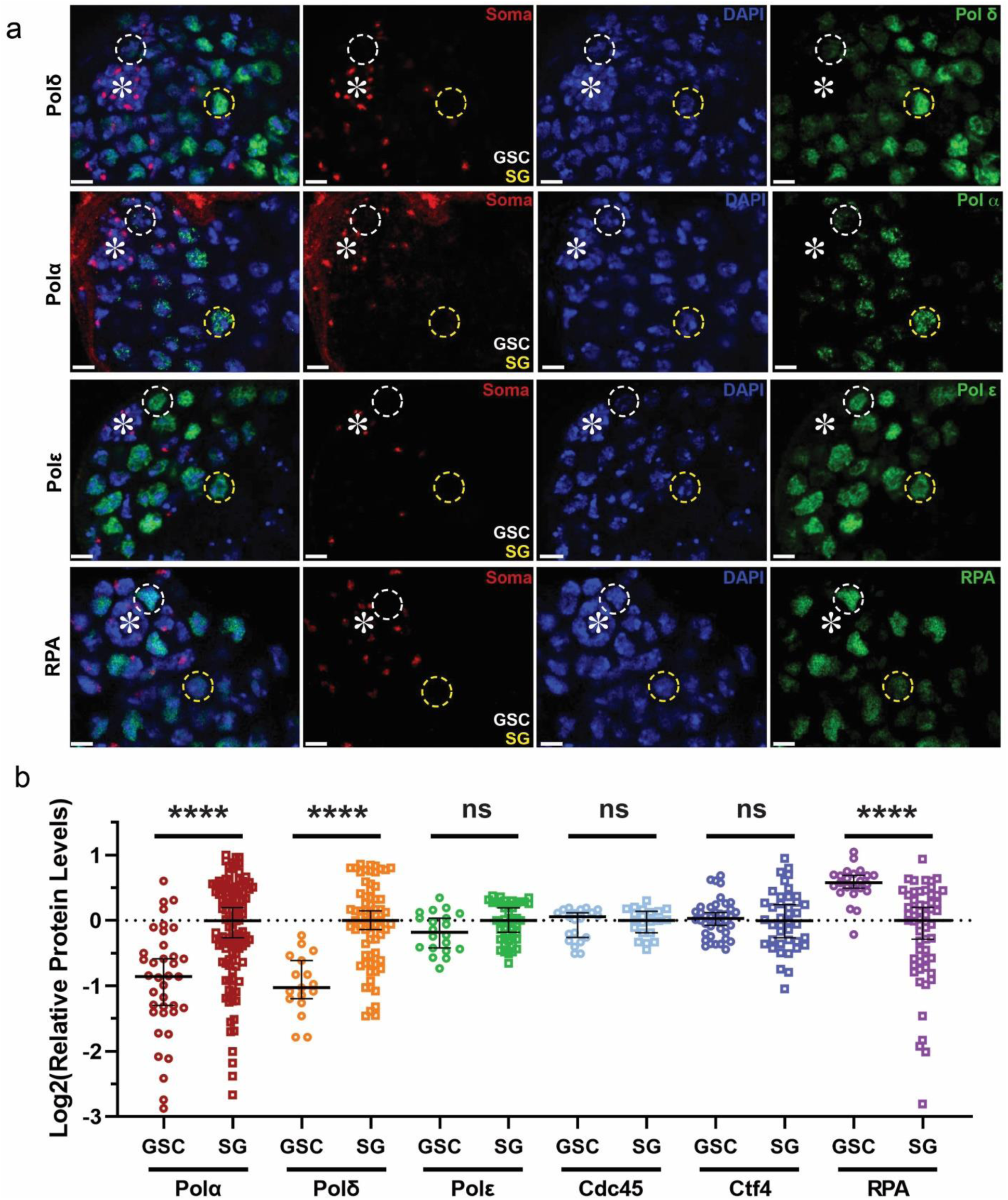
Distinct expression patterns of different replication machinery components in the *Drosophila* male germline stem cell lineage. (**a**) Images of expression of 3×HA tagged endogenous DNA polymerases Polδ, Polα, Polε (see Materials and Methods), as well as the RPA 70-EGFP expressed from a transgene with its own promoter (*44*). Somatic cell-enriched histone modification H4K20me2/3 (red) (*91*), DAPI (blue), and the respective replication proteins (green). Both Polδ and Polα show decreased levels in GSCs, while Polε has comparable expression between GSCs and SGs. Contrastingly, RPA is more enriched in GSCs compared to SGs using a transgene under the control of its endogenous regulatory elements (*rpa70>rpa70-EGFP*) (*44*). Representative GSCs are indicated by the white dotted circle while SGs are indicated by the yellow dotted circle. (**b**) Quantification of the relative expression levels of different replication proteins, using a batch-based normalization to GSCs from the corresponding testis sample followed by log_2_ transformation (Materials and Methods). Medians: GSC Polα log_2_= −0.85 (n= 37), SG Polα log_2_= 0.00 (n= 118); GSC Polδ log_2_= −1.03 (n= 17), SG Polδ log_2_= 0.00 (n= 70); GSC Polε log_2_= −0.18 (n= 20), SG Polε log_2_= 0.00 (n= 43); GSC Cdc45 log_2_= 0.06 (n= 21), SG Cdc45 log_2_= 0.00 (n= 21); GSC Ctf4 log_2_= 0.03 (n= 40), SG Ctf4 log_2_= 0.00 (n= 40); GSC RPA log_2_= 0.58 (n= 23), SG RPA log_2_= 0.00 (n= 57). Asterisk: hub. Scale bars: 10 μm. All ratios: Median ± 95% Confidence Interval (CI). Mann-Whitney test, ****: *P*< 10^-4^, ns: not significant. See Table S1 for details.

Quantification of these results reveals that GSCs have approximately 51% the levels of Polδ and 58% the levels of Polα as compared to SGs, while Polε levels are comparable between GSCs and SGs (Fig. 1a, b). In contrast, RPA is 1.54-fold more enriched in GSCs than in SGs (Fig. 1a, b). However, other replication machinery components, such as the replication fork progression Cell Division Cycle protein 45 (Cdc45), show no significant difference between GSCs and SGs (Fig. 1b). Another component whose yeast homolog has been shown to have histone chaperoning activities (*22*), Chromosome Transmission Fidelity 4 (Ctf4), also displays similar levels between GSCs and SGs (Fig. 1b, Fig. S1a).

The significantly reduced levels of lagging strand polymerases in GSCs could lead to relatively delayed lagging strand synthesis, which could result in excessive ssDNA. The higher levels of RPA in GSCs could be responsible for coating and stabilizing ssDNA (*45*). Moreover, RPA is capable of competing with Polα at ssDNA sites, therefore preventing Polα from binding to and acting on the lagging strand (*46–50*). Therefore, decreased Polα and increased RPA could cooperatively contribute to measured lagging strand synthesis in the GSCs, which could also underlie the longer cell cycle length of GSCs than SGs, as reported previously (*51*).

### Reducing Polα levels or inhibiting Polα activities increase old *versus* new histone separation in S-phase nuclei of progenitor cells

Based on the above observation, we hypothesize that relatively slow lagging strand synthesis could bias old histone recycling to the finished leading strand at individual replication forks, serving as a key molecular mechanism underlying asymmetric histone incorporation in GSCs. To investigate this hypothesis, we first examined the distribution of old *versus* new histones in intact nuclei using a dual-color system to label old H3 with EGFP and new H3 with mCherry in the male germline (*15, 19*). To avoid any possible complications caused by non-chromatin-bound histones, we used a stringent clearance buffer that has been shown to effectively remove free histones in the nucleus (*19, 52, 53*). This strategy, together with high spatial resolution Airyscan microscopy (*15, 54*), allows us to visualize separable old H3-EGFP *versus* new H3-mCherry enriched regions in the control *wild-type* (WT) GSCs at S-phase, labeled with a pulse of thymidine analog 5-ethynyl-2’-deoxyuridine (EdU) (Fig. 2a, b). In contrast, the degree of separation between old and new H3 is less in WT SGs than that in WT GSCs during S-phase (Fig. 2b). Quantification using a relative Pearson colocalization measurement (*16, 55, 56*) reveals significantly higher degree of colocalization between old H3-EGFP and new H3-mCherry in WT SGs than in WT GSCs (Fig. 2c), consistent with asymmetric incorporation of old *versus* new histones in S-phase WT GSCs.

**Figure 2:**
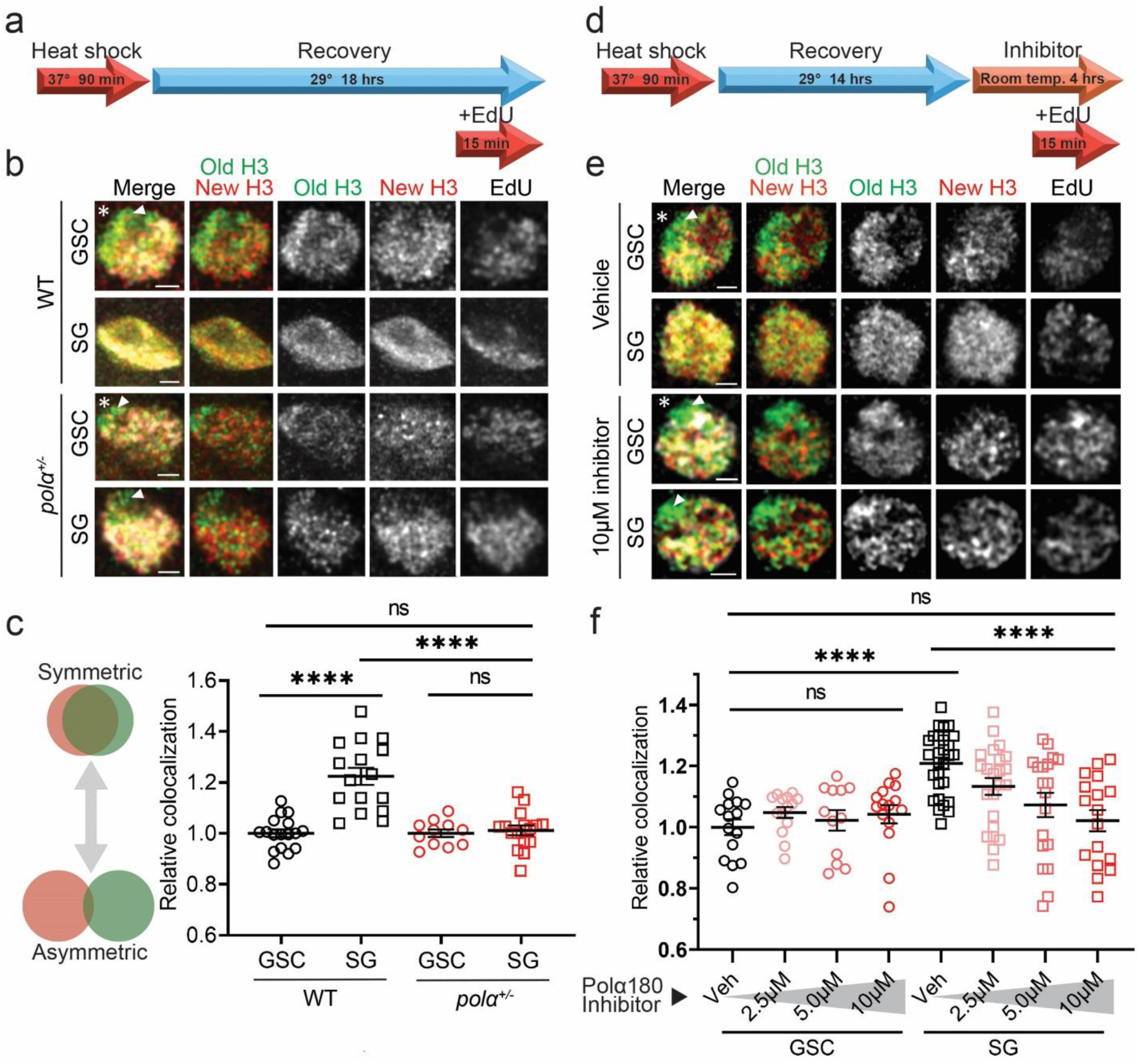
Reducing Polα levels or compromising Polα activities increases old histone *versus* new histone separation in S-phase nuclei of progenitor cells. (**a**) Regime testing old (EGFP) vs. new (mCherry) histone localization pattern following heat shock-induced tag switch. (**b**) Airyscan images of representative control *wild-type* (WT) GSC, WT SG, *polα50^+/-^* GSC, and *polα50^+/-^* SG, respectively, in S-phase nuclei wherein nucleoplasmic histones are largely washed off using a stringent clearance buffer. In all merged images: old H3 (green), new H3 (red), as well as EdU (white), Arm (not shown but used as hub marker). Asterisk: hub. Scale bars: 1 μm. (**c**) Quantification of the correlation between old H3 and new H3 signals in S-phase nuclei using a batch-based normalization to control GSCs (Materials and Methods): WT GSC= 1.00± 0.06 (n= 17), WT SG= 1.90± 0.13 (n= 16), *polα50^+/-^* GSC= 1.00± 0.06 (n= 11), *polα50^+/-^* SG= 1.05± 0.08 (n= 16). See Table S3 for details. All images and quantifications for SGs use the 4-cell SGs. (**d**) Regime testing old vs. new histone localization pattern in response to Polα180 (or Pol A1) inhibitor. (**e**) Airyscan images of representative GSCs and SGs treated with vehicle or Polα180 inhibitor for four hours prior to clearance buffer treatment and fixation. Arrowheads in (**b**) and (**e**): unreplicated regions are enriched with old H3 but depleted with new H3 and EdU labeling. (**f**) Quantification of the correlation between old H3 and new H3 signals in S-phase nuclei following inhibitor treatment using a batch-based normalization to vehicle-treated GSCs (Materials and Methods): Vehicle GSC= 1.00 ± 0.03 (n= 15), 2.5μM GSC= 1.05 ± 0.02 (n= 14), 5.0μM GSC= 1.02 ± 0.03 (n= 12), 10μM GSC= 1.04 ± 0.03 (n= 15); Vehicle SG= 1.21 ± 0.02 (n= 28), 2.5μM SG= 1.13 ± 0.03 (n= 23), 5.0μM SG= 1.07 ± 0.04 (n= 19), 10μM SG= 1.02 ± 0.04 (n= 17). See Table S4 for details. All images and quantifications for SGs use the 4-cell SGs. All ratios: Mean ± Standard Error of the Mean (SEM). Mann-Whitney test, ****: *P*< 10^-4^, ns: not significant.

We next asked whether compromising lagging strand synthesis in SGs could recapitulate GSC-like features, such as separable old *versus* new histones in the S-phase nuclei. Since replication components are essential for animal survival and cell cycle progression, we sought to compromise lagging strand polymerases without causing cell cycle arrest or cell death, which does occur in strong loss-of-function homozygous or in RNAi knockdown germ cells (data not shown). However, by using a null allele of the *polα50* gene (Materials and Methods), which encodes the DNA Primase Subunit 1 (or Prim1), we generated *polα50^+/-^* heterozygotes. We were able to maintain viable flies with no detectable systematic phenotypes. Intriguingly, when the Primase levels are reduced in *polα50^+/-^* males, S-phase SGs display much more separable patterns between old and new H3, to a level indistinguishable from WT GSCs as well as *polα50^+/-^*GSCs (Fig. 2b, c).

In addition to this genetic approach, we tried a pharmacological strategy with a Polα inhibitor that prevents the DNA binding ability and primer elongation activity of DNA Polymerase α subunit 1 (PolA1 or Polα180, Fig. S2a) (*57*). At a high concentration (e.g., 100μM), this inhibitor completely blocks DNA replication, indicated by the absence of EdU incorporation in different staged germ cells (data not shown). However, when using this inhibitor at a relatively low concentration (e.g., 10μM), normal DNA replication could proceed with proper EdU incorporation compared to the control sample treated with vehicle (Fig. 2d, Fig. S2b-c). With this inhibitor treatment, SGs exhibit separable old *versus* new H3 patterns similar to those detected in the *polα50^+/-^*cells (inhibitor SG in Fig. 2e vs. *polα50^+/-^* SG in Fig. 2b). The inhibitor-treated SGs show more separation than the vehicle-treated SGs, and display patterns comparable to either inhibitor-treated GSCs or vehicle-treated GSCs (Fig. 2e). Quantifications further reveal that this inhibitor induces old *versus* new H3 separation in SGs in a dosage-dependent manner, but it causes insignificant changes in GSCs regardless of the concentration (Fig. 2f). Notably, the presence of intermediate histone separation patterns at decreasing concentrations of inhibitor (e.g., 5.0μM and 2.5μM) indicate that the asymmetric histone incorporation pattern is tunable and scales to the inhibition of Polα. Additionally, because GSCs are relatively unaffected, we hypothesize that these cells are at the maximum of histone asymmetry and thus cannot be made more asymmetric by compromising Polα.

Moreover, in all imaged nuclei undergoing DNA synthesis, unreplicated regions are enriched with old H3 but devoid of new H3 as well as EdU labeling (arrowheads in Fig. 2b and Fig. 2e), confirming that actively replicating regions are coupled with new H3 incorporation. Together, these data in intact S-phase nuclei demonstrate that reducing primase levels or inhibiting Polα activity are each sufficient to induce separable old *versus* new histone incorporation in S-phase SGs, to a degree indistinguishable from that in GSCs.

### Reducing Polα levels enhances asymmetric old histone incorporation at the replication fork in S-phase progenitor cells

Next, in order to directly visualize the dynamic histone incorporation patterns at the actively replicating regions, a short pulse of EdU was introduced in combination with a single-molecule chromatin fiber technique (*15, 17*). To precisely label chromatin fibers derived from GSCs *versus* non-stem progenitor SGs, we paired the Gal4 transcription activator controlled by the early germline-specific *nanos* driver (*nos-Gal4ΔVP16*) (*58*) with the Gal80 transcription repressor under the control of the *bag of marbles* promoter (*bam-Gal80*), which turns on expression from 2-cell to late stage SGs (*59*). This combination restricts the *H3-EGFP* transgene expression almost exclusively in GSCs with some detectable expression in the gonialblasts (GBs) but almost undetectable signals in the SGs (Fig. S1d), which differs from the early-stage germ cell expression pattern driven solely by *nos-Gal4* (*60*) (Fig. S1b) and late-stage germ cell expression pattern driven solely by *bam-Gal4* (*61–63*) (Fig. S1c). These germline stage-specific expression patterns are confirmed by quantification using a *H3-EGFP* reporter (Fig. S1e).

Using the *H3-EGFP* reporter with different drivers, we labeled chromatin fibers derived from early-stage germ cells including GSCs (*nos>H3-EGFP*), from very early-stage germ cells enriched with almost exclusive GSCs (*nos-Gal4ΔVP16; bam-Gal80*>*H3-EGFP*), and from late-stage SGs (*bam*>*H3-EGFP*). We then explored old histone recycling patterns at the H3-EGFP-labeled and EdU-positive chromatin fibers using the old H3-enriched H3K27me3 histone modification (*31, 64, 65*). We also distinguished the strandedness with the lagging strand-enriched component Proliferating Cell Nuclear Antigen (PCNA) (*15, 66*). Together, chromatin fibers carrying all four markers (i.e., H3-EGFP, EdU, anti-H3K27me3, and anti-PCNA) were analyzed using super-resolution Airyscan microscopy. While *nos>H3-EGFP*-labeled chromatin fibers show a relatively wide distribution of H3K27me3 between replicative sister chromatids (Fig. S3a, b) with an overall biased distribution toward the PCNA-depleted leading strand (Fig. 3f, Fig. S3c), *nos-Gal4ΔVP16; bam-Gal80*>*H3-EGFP*-labeled chromatin fibers show consistently more asymmetric H3K27me3 distribution toward the leading strand (Fig. 3a, f, Fig. S3c). In contrast, *bam*>*H3-eGFP*-labeled fibers display a more symmetric H3K27me3 distribution pattern (Fig. 3b, f, Fig. S3c). Notably, previous reports using an imaging-based proximity ligation assay in intact nuclei (*67, 68*) demonstrate that new histones have a substantial lagging strand preference in GSCs but not in SGs (*15*), consistent with the results shown here.

**Figure 3:**
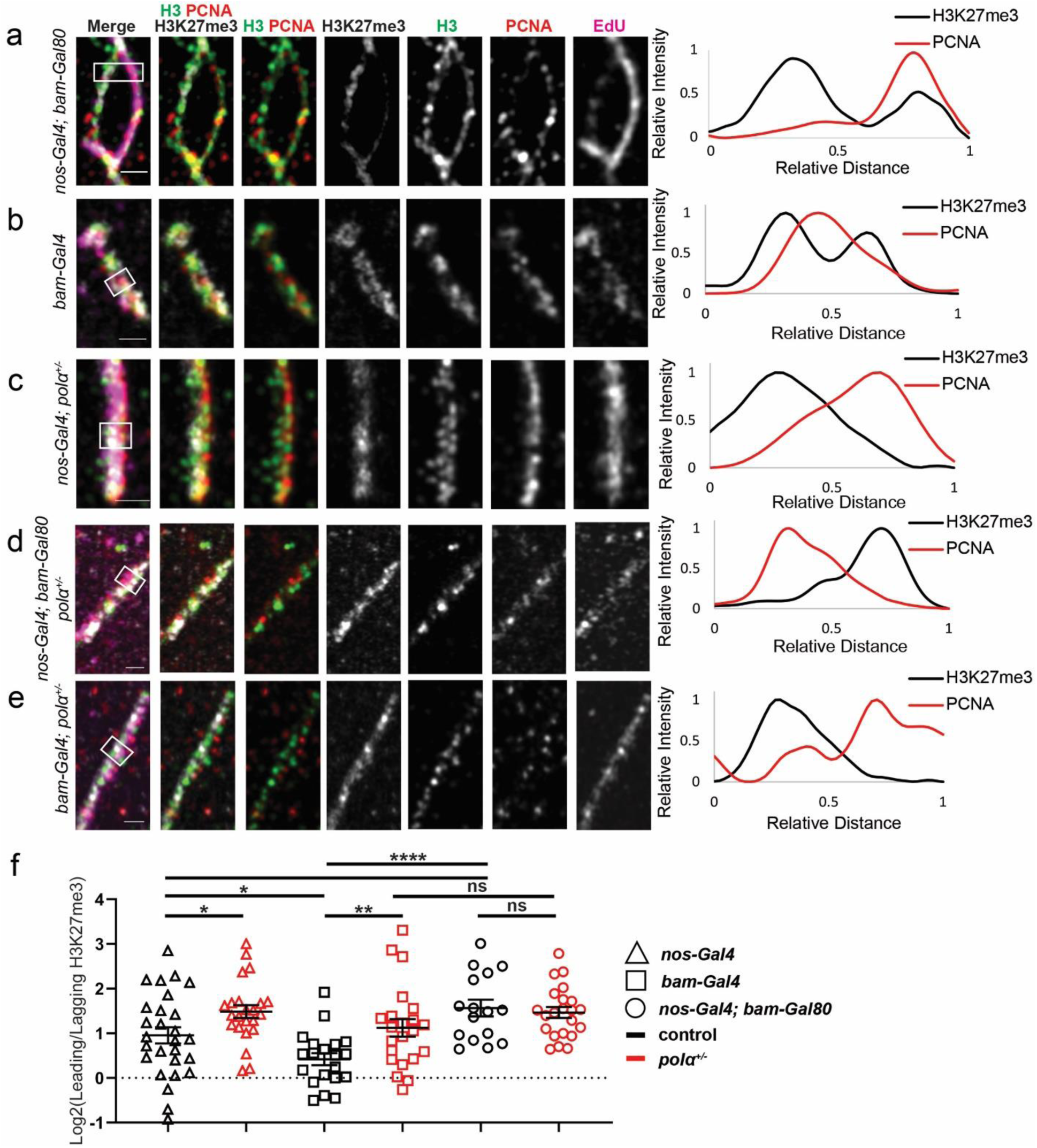
Reducing Polα levels enhances asymmetric old histone recycling at the replication fork in progenitor cells. (**a-e**) Airyscan images of chromatin fibers isolated from testes with the following genotypes: (**a**) *nos-Gal4ΔVP16; bam-Gal80>H3-EGFP*, (**b**) *bam-Gal4>H3-EGFP*, (**c**) *nanos-Gal4>H3-EGFP; polα50^+/-^*, (**d**) *nos-Gal4ΔVP16; bam-Gal80>H3-EGFP; polα50^+/-^*, (**e**) *bam-Gal4>H3-EGFP; polα50^+/-^*, respectively. In all merged images: H3K27me3 (white), H3-EGFP (green), PCNA (red), and EdU (magenta). All images are accompanied by a line plot showing the distance-dependent H3K27me3 and PCNA signals over the indicated region (white outlined box). (**f**) Quantification of the H3K27me3 signals on chromatin fibers in log_2_ scale: *nanos-Gal4>H3-EGFP*= 0.95± 0.18 (n= 27), *nanos-Gal4>H3-EGFP; polα50^+/-^*= 1.49± 0.15 (n= 23), *bam-Gal4>H3-EGFP*= 0.42± 0.14 (n= 20), *bam-Gal4>H3-EGFP; polα50^+/-^*= 1.12± 0.20 (n= 22), *nos-Gal4ΔVP16; bam-Gal80>H3-*EGFP= 1.56± 0.19 (n= 16), *nos-Gal4ΔVP16; bam-Gal80>H3-EGFP; polα50^+/-^*= 1.66± 0.15 (n= 22). Scale bars: 1 μm. All ratios: Mean± SEM. Mann-Whitney test, ****: *P*< 10^-4^, **: *P*< 0.01, *: *P*< 0.05, ns: not significant. See Table S6 for details.

To quantify old histone incorporation patterns, we used H3K27me3 as a proxy for old histones and plotted its ratio on the PCNA-depleted leading strand to the PCNA-enriched lagging strand (log_2_ ratios in Fig. 3f, Fig. S3c-d). The *nos-Gal4ΔVP16; bam-Gal80*-labeled and the *bam*-labeled chromatin fibers are not only statistically distinguishable from each other (*P*< 10^-4^, Fig. 3f), but also statistically different from the *nos*-labeled group (*P*< 0.05, Fig. 3f). Interestingly, combining the *nos-Gal4ΔVP16; bam-Gal80*-labeled and *bam*-labeled groups *in silico* generates a data set indistinguishable from the *nos*-labeled group (Fig. S3c), suggesting that the heterogeneity of both GSC-derived and SG-derived fibers could underlie the detected H3K27me3 variation among the *nos*-labeled chromatin fibers.

Furthermore, we found that the *nos*-labeled fibers from *polα50^+/-^*testes show more asymmetric H3K27me3 distribution toward the leading strand than the *nos*-labeled fibers from the control (Fig. 3c, f). Consistently, the *nos*-labeled fibers from heterozygotes of the *polα180* gene, which encodes DNA Polymerase α subunit 1 (or PolA1), also exhibit a more asymmetric H3K27me3 distribution pattern toward the leading strand than those from the control (Fig. S3d-e). In contrast, compromising Polα50 has little effect on *nos-Gal4ΔVP16; bam-Gal80*-labeled chromatin fibers (Fig 3d, f). Notably, *bam>H3-eGFP*-labeled chromatin fibers display significantly more asymmetric patterns in the *polα50^+/-^* samples than in the control (Fig 3e, f), in accordance with the results shown in intact S-phase nuclei (Fig. 2b, c). Together, these results demonstrate that compromising Polα affects SGs with normally high levels of Polα (i.e., *bam*-labeled chromatin fibers in Fig. 3 and intact SG nuclei in Fig. 2) more than GSCs that already have low levels of Polα (i.e., *nos-Gal4ΔVP16; bam-Gal80*-labeled chromatin fibers in Fig. 3 and intact GSC nuclei in Fig. 2). It is likely that reducing Polα levels below a certain threshold cannot further increase histone asymmetry, but reducing Polα from relatively high levels (i.e., SG-like) to relatively low levels (i.e., GSC-like) is sufficient to enhance asymmetric old histone recycling at the replication fork.

Finally, to test whether RPA also contributes to asymmetric histone incorporation, we overexpressed the *rpa70* cDNA using *nos-Gal4* (*nos>rpa70-HA)*. Likewise, the overexpression of RPA70 results in enhanced asymmetric H3K27me3 incorporation at the replicative regions, indicating that increased levels of RPA lead to enhanced asymmetric old histone recycling (Fig. S3d, f). Notably, these results are consistent with the previous report that using the *rpa-70>rpa-70-EGFP* line, where the transgenic RPA-70-EGFP fusion protein is under the control of the endogenous *rpa-70* regulatory elements and represents a slight overexpression condition. Under this condition, an average of 3.2-fold leading strand biased H3K27me3 asymmetry is detected, more than the control line which shows an average of 2.0-fold leading strand biased H3K27me3 asymmetry (*15*). These effects could be attributed to the previously reported competing roles of RPA in preventing Polα from binding to the lagging strand (*46–50*). In summary, the chromatin fiber results demonstrate that SGs with relatively high levels of Polα and low levels of RPA can be induced to have GSC-like asymmetric old histone incorporation at the replicative regions by reducing Polα levels or by enhancing RPA expression.

### Reducing Polα levels induces differential condensation of old histone-*versus* new histone-enriched regions in M-phase progenitor cells

It has been reported that old H3-*versus* new H3-enriched chromosomal regions display differential condensation in the M-phase GSCs but overlapping pattern in the M-phase SGs (*19*). Consistent with previous reports (*19, 53*), the control GSCs and SGs display marked condensation differences between old H3-and new H3-enriched regions (Fig. 4a, b), while the *polα50^+/-^* SGs (Fig. 4d) show GSC-like (Fig. 4a, c) differential condensation patterns. Here, using a relative chromatin compaction index to measure the differential condensation between old H3-and new H3-enriched regions as reported previously (*19, 53*), significant difference could be detected between GSCs and SGs in the control testes but not between GSCs and SGs in the *polα50^+/-^* testes (Fig. 4e). Importantly, in the *polα50^+/-^*testes, both GSCs and SGs display similar patterns compared to the control GSCs but significantly distinct patterns compared to the control SGs (Fig. 4e). Collectively, these results demonstrate that by compromising a single lagging strand-enriched component, differential condensation of old H3-*versus* new H3-enriched regions in M-phase cells, a GSC-specific feature, can be recapitulated in the SGs.

**Figure 4:**
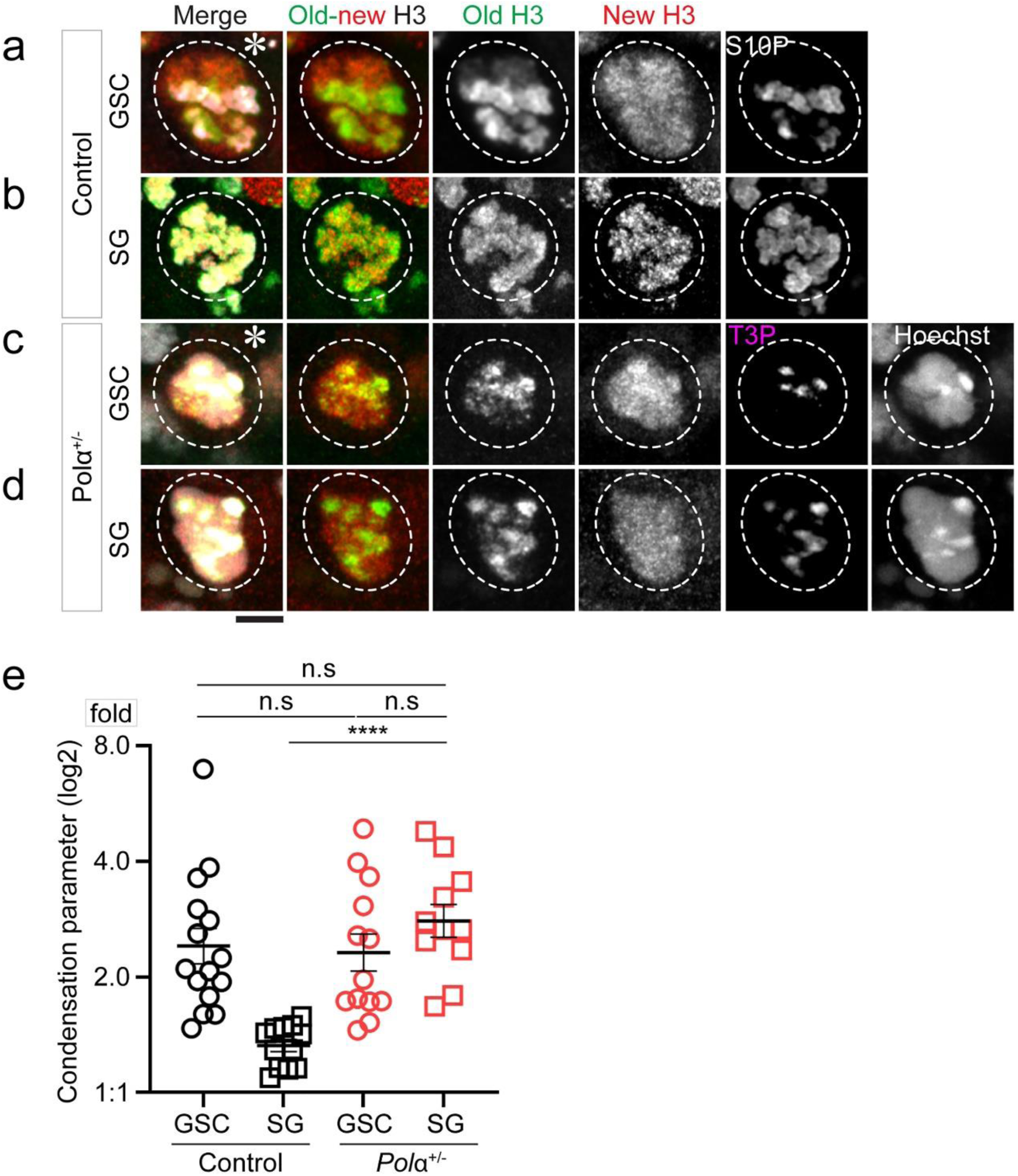
Reducing Polα levels induces differential condensation of old H3 *versus* new H3 enriched regions in M-phase progenitor cells. (**a-b**) Representative images of: (**a**) an M-phase GSC showing more compact old H3-enriched regions than new H3-enriched regions (positive with a mitotic marker anti-H3S10ph, H3S10P or S10P (*19*); (**b**) an M-phase 8-cell SG showing equally compact old H3-enriched and new H3-enriched regions (positive with S10P) in the control *wild-type* testes. (**c-d**) Representative images of: (**c**) an M-phase GSC and (**d**) an M-phase 8-cell SG in the *pola50^+/-^* testes, both showing more compact old H3-enriched regions than new H3-enriched regions (positive with a mitotic marker anti-H3T3ph, H3T3P or T3P(*92*). (**e**) Compaction index in log_2_ scale: Control GSC= 1.27± 0.15 (n=15), Control 8-cell SG= 0.41± 0.05 (n=12), *polα50^+/-^* GSC= 1.21± 0.15 (n=13), and *polα50^+/-^* 8-cell SG= 1.48± 0.14 (n=11). The control compaction index data are from (*19*) with permission. See Table S8 for details. All ratios: Mean± SEM. Mann-Whitney test, ****: *P*< 10^-4^, ns: not significant.

### Detectable asynchrony between leading strand and lagging strand syntheses

Next, to measure the leading *versus* lagging strand syntheses in the early-stage germline, we attempted to directly visualize these processes using active incorporation of nucleotide analogs. Previously, it has been shown that the syntheses of the two DNA strands can be discontinuous where the leading and lagging strand polymerases are not tightly coupled in *E. coli* (*69*) or when applying PolA1 inhibitor in cultured human cells (*70*). Recently, it has been reported that the temporal differences in replicating leading strand *versus* lagging strand biases old histone incorporation by the strand more closely coupled to the replication fork progression in *S. cerevisiae* (*71*). To investigate whether this temporal difference exists and is detectable in the *Drosophila* testes, we investigated whether leading strand *versus* lagging strand syntheses can be differentially labeled, using distinct nucleotide analogs introduced in a sequential order [e.g., a short pulse of EdU followed by a short pulse of Bromodeoxyuridine (BrdU), Fig. 5a]. Using this regime, DNA fibers where both strands are co-labeled with just one nucleotide (e.g., EdU) should represent regions where both strands are replicated within the time window of the EdU pulse (an example is shown in the top panel of Fig. 5b). However, DNA fibers with EdU and BrdU on opposing strands likely capture the uncoupled syntheses of the two strands (an example is shown in the bottom panel of Fig. 5b). Indeed, the DNA fibers derived from the apical testis tips display the latter pattern in approximately 40% of the fibers (Fig. S4a). On average, DNA fibers carrying both EdU and BrdU display a 2.35-fold BrdU enrichment toward one strand whereas a 1.91-fold EdU enrichment toward the opposing strand (Fig. 5c, Fig. S4b).

**Figure 5:**
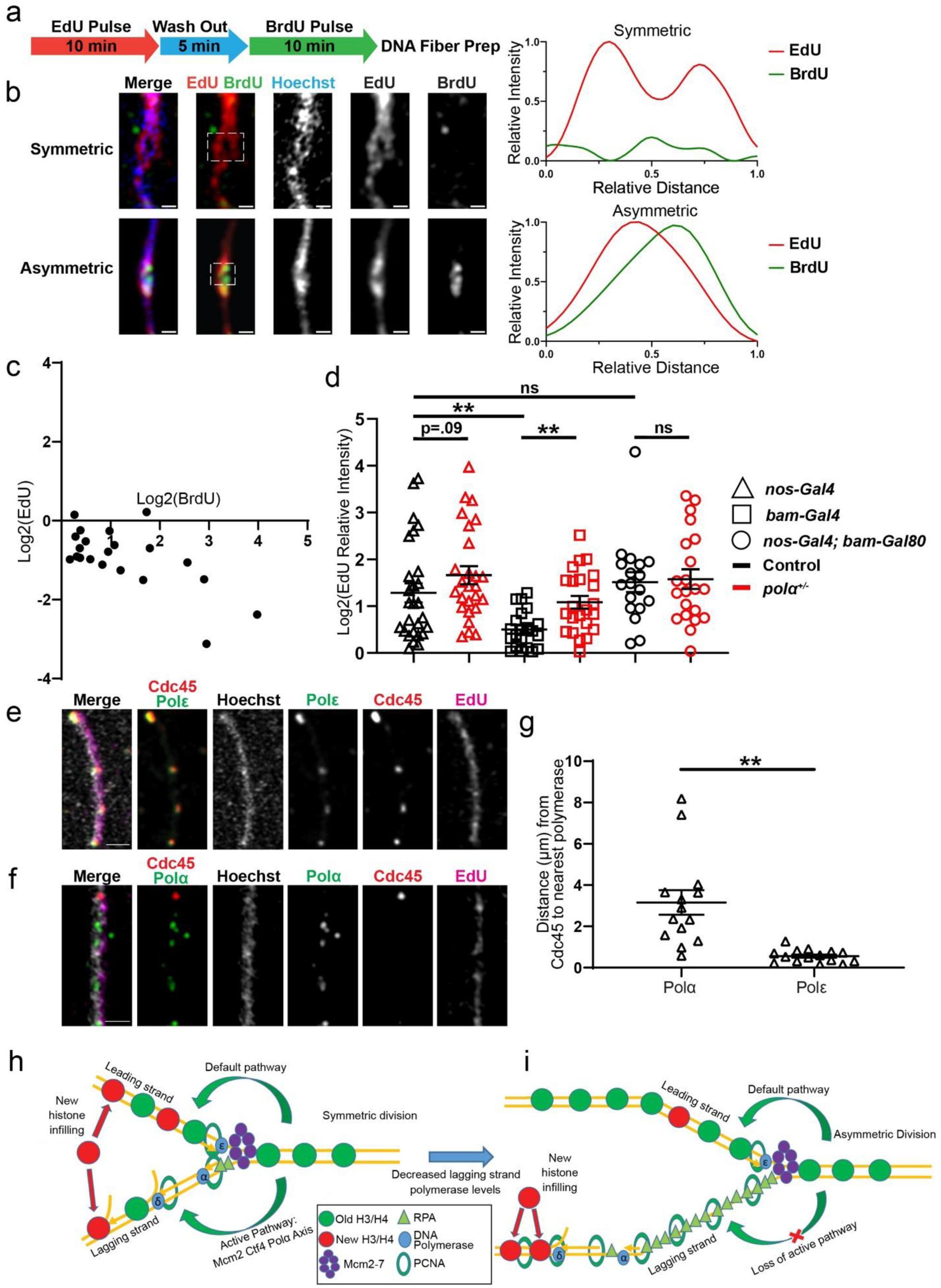
Asynchronous leading strand *versus* lagging strand syntheses. (**a**) Regime with a 10-min EdU pulse followed by a 5-min wash out, then a 10-min BrdU pulse to label DNA fibers (Materials and Methods) (**b**) Airyscan images of DNA fibers: The two line-plots correspond to a representative symmetric replicative region with EdU labeling both strands (white dotted outline on the top panel) and an asymmetric region where EdU and BrdU are on the opposing strands (white dotted outline on the bottom panel). (**c**) Log_2_-scale 2D plot shows the distribution of BrdU and EdU on the DNA fibers with both signals. Most fibers display EdU and BrdU on the opposing DNA strands. See Table S10 for details. (**d**) Quantification of EdU distribution in log_2_-scale, introduced by a 15-minute EdU pulse labeling, on chromatin fibers labeled with H3-EGFP driven by the following drivers without strandedness information: *nanos-Gal4*= 1.29± 0.21 (n= 27), *nanos-Gal4; polα50^+/-^*= 1.66± 0.19 (n= 26), *bam-Gal4*= 0.50± 0.09 (n= 21), *bam-Gal4; polα50^+/-^*= 1.08± 0.14 (n= 23), *nos-Gal4ΔVP16; bam-Gal80*= 1.52± 0.21 (n= 18), *nos-Gal4ΔVP16; bam-Gal80; polα50^+/-^*= 1.58± 0.21 (n= 21). Scale bars: 1 μm. All ratios: Mean± SEM. Mann-Whitney test, **: *P*< 0.01, ns: not significant. See Table S11 for details. (**e-f**) Visualization of delayed lagging strand synthesis: Airyscan image of representative chromatin fibers labeled with endogenous Cdc45-mcherry (red), EdU (magenta), Hoechst (white), along with (**e**) Polε-HA (green) or (**f**) Polα-HA (green). (**g**) Quantification of the distance along chromatin fibers from the center of the Cdc45 signal to the nearest signal of the DNA polymerase: Polα= 3.16± 0.60 μm (n= 14), Polε= 0.56± 0.08 μm (n= 15). Scale bars: 1 μm. All ratios: Mean± SEM. Mann-Whitney test, ****: *P*< 10^-4^. See Table S12 for details. (**h-i**) Models depicting how reducing lagging strand polymerase levels could drive increased histone asymmetry at the replication fork: (**h**) In symmetrically dividing cells, comparable leading strand *versus* lagging strand syntheses give the old histone equal opportunities to be recycled by either strand, based on previous reports(*21, 22, 26*). (**i**) In asymmetrically dividing cells, reduced levels of lagging strand polymerases lead to measured lagging strand synthesis relative to the leading strand, which results in a temporal difference and biases old histone recycling by the leading strand, whereas new histones infill to the lagging strand.

On the other hand, when a single short EdU pulse is introduced on early germline-derived chromatin fibers where strandedness can be determined, 52% of them display a strong bias (>2-fold) toward one of the two strands, with 79% of them displaying strong asymmetry toward the lagging strand (Fig. S4c-d), likely due to a longer time for synthesizing the lagging strand and thus a higher opportunity for the lagging strand to be labeled (Fig. S4e). Consistently, PCNA signals often display asymmetric distribution on early germline-derived chromatin fibers, along with EdU, toward the H3K27me3-depleted lagging strand (Fig. S4d, Fig. 3a, and Fig. S3a).

To further test whether delayed lagging strand synthesis is mechanistically responsible for asymmetric histone incorporation, we analyzed the EdU signals of all chromatin fibers that have been analyzed for the H3K27me3 patterns in Figure 3. Indeed, very early-stage germline-derived (*nanos-Gal4ΔVP16; bam-Gal80*) chromatin fibers show a high degree of asymmetric EdU patterns, while the late-stage germline-derived (*bam-Gal4*) chromatin fibers primarily show symmetric EdU distribution between sister chromatids (Fig. 5d, Fig. S4f). Notably, EdU asymmetry is substantially enhanced in the late-stage germline-derived (*bam-Gal4*) chromatin fibers from the *polα50^+/-^*testes than that from the control (*P*< 0.01 in Fig. 5d, Fig. S4f). The small incidence of leading strand-but a large population of lagging strand-biased EdU incorporation (Fig. S4d, f) are consistent with the temporal asynchrony between leading strand and lagging strand syntheses, with the lagging side synthesizing DNA more slowly on average (Fig. S4e). Overall, these results show that compromising Polα is sufficient to increase asymmetric H3K27me3 incorporation by the leading strand, likely by enhancing the temporal asynchrony between sister chromatid syntheses.

Finally, to directly visualize delayed lagging strand synthesis, we used an endogenously tagged *cdc45* gene, resulting in the Cdc45-mCherry fusion protein as a marker for the Cdc45-MCM-GINS (CMG) complexes (*72, 73*), in order to label actively progressing replication forks. We then performed immunostaining using antibodies against the HA tag to label Polα-HA for lagging strand polymerase and Polε-HA for leading strand polymerase, using the endogenously tagged genes, respectively. Remarkably, while Polε is always tightly associated with Cdc45 at EdU-labeled replicative chromatin fibers (Fig. 5e, g), Polα could be found in tracts extending away from Cdc45 (Fig. 5f, g), indicating that Polα is spatially decoupled from the actively progressing fork. Together, these results demonstrate that the temporal and spatial separation between leading strand and lagging strand syntheses can be visualized by nucleotide analogs introduced at different time points during DNA replication or with different replisome components.

## Discussion

Here, we report a crucial molecular mechanism underlying asymmetric histone incorporation in stem cells. In symmetrically dividing progenitor cells, comparable leading *versus* lagging strand syntheses give the old histone equal opportunities to be recycled by both strands (Fig. 5h), as shown previously (*21, 22, 26, 28, 31*). In asymmetrically dividing germline stem cells, reduced lagging strand polymerase levels could slow down lagging strand synthesis relative to the leading strand, which results in a pronounced temporal difference. This difference could bias displaced old histone ahead of the fork to be immediately recycled by the leading strand whereas new histones infill to the lagging strand (Fig. 5i). This model is consistent with a previous report that nucleosomes have the priority to be reincorporated by the double-stranded leading strand *in vitro* (*74*). The increased expression of RPA in stem cells could also facilitate this process (Fig. 5i). This model is consistent with the previous imaging-based results displaying abundant RPA bound to the lagging strand on early-stage germ cell derived chromatin fibers (*15*). Intriguingly, either reducing the expression of the key lagging strand polymerase or inhibiting its activity is sufficient to induce stem cell-specific asymmetric histone incorporation patterns even in non-stem progenitor cells.

Notably, even though SGs with approximately 50% Primase (i.e., *pola50^+/-^*) have S-phase replication-dependent histone incorporation patterns and M-phase differential chromosomal condensation patterns similar to those in GSCs, SGs do not reside in a polarized microenvironment like the “niche” for GSCs. Additionally, there is no evidence that the microtubule organization centers, the centrosomes, have asymemtric activities in the *wild-type* SGs (*62, 75*). Therefore, these chromatin asymmetries may not result in substantial differences between the two daughter cells resulting from SG symmetric cell division, unlike the asymmetric division of GSCs. Finally, the *pola50^+/-^* SGs seem to undergo terminal differentiation properly, as there are no obvious germline defects detectable in the *pola50^+/-^*males. It is plausible that the molecular features such as the transcriptome of the *pola50^+/-^* SGs remain unchanged or have inconsequential changes, despite the detectable changes of their chromatin structure. This indicates that the epigenome potentiates cell fate change but may not be determinstic for such a decision. Remarkably, the 50% reduction of Primase in heterozygous SGs is analogous to the protein level change detected in WT GSCs (Fig. 1b). Further reduction of Primase will cause cell cycle arrest, contributing to severe defects detected in homozygotes and under significant knockdown conditions (data not shown).

Furthermore, we focus on the Polα-primase complex in this study because it predominantly acts on the lagging strand except the initial priming event on the leading strand and during rare re-priming events at stalled replication forks (*47, 76, 77*). On the other hand, Polδ could contribute to the synthesis of both the leading strand and the lagging strand (*78, 79*). Although it is known that Polα (*22, 28*) and RPA (*80*) also play a role in chaperoning histones during replication-coupled nucleosome assembly, such activities have been demonstrated using specific mutations at their histone interacting domains. Here, our studies make use of genetic approaches that either compromise function or change expression of the entire proteins. Additionally, the pharmacological method utilizes enzyme inhibitors whose effects are more related to their roles as replication machinery components, such as primer elongation, rather than chaperones. Notably, DNA replication inhibitors are often used to target over-proliferative cancer cells. Indeed, the drug adarotene and its derivative molecule used in this study have been shown to have anti-cancer properties in mice (*57, 81, 82*). However, the dose used in our studies is much lower than the dose used for cancer therapy to ensure minimal effect on S-phase progression (Fig. S2b-c). Therefore, both genetic and pharmacological approaches emphasize the importance of the control for the optimal Polα level or inhibitor dosage, which needs to be calibrated empirically in different systems.

Finally, DNA replication is fundamentally an inherently asymmetric process wherein the synthesizing processes of the leading strand *versus* the lagging strand are widely divergent. Previous studies have shown examples of uncoupled leading strand *versus* lagging strand syntheses in bacteria and cultured cells (*69–71, 83*), particularly in cases where Polα or its priming activity is compromised. It has long been recognized that the leading strand *versus* the lagging strand may have the potential to differentially incorporate nucleosomes (*84*).

Intriguingly, the old histone-enriched H3K9me3 has been shown to be recycled by the leading strand at the retrotransposon elements in order to repress their ectopic transcription in S-phase mouse embryonic stem cells (*85*). Further, it has been shown that DNA replication speed and timing underlie cell fate regulation in mammalian cells, including mouse and human cells (*86–89*). Here, our results indicate that the inherent asymmetry of DNA replication itself could be utilized to differentially regulate histone incorporation and this process displays stage specificity within an endogenous adult stem cell lineage. These results point to a very exciting possibility that developmentally programmed expression of key DNA replication components could regulate the establishment of distinct epigenomes in a cell type-and stage-specific manner. Given that replication components as well as histone proteins and their respective modifications are highly conserved, exploring how this mechanism may be utilized in other developmental contexts across different multicellular organisms could be a very intriguing research direction (*5, 90*). These elegant and efficient mechanisms could be used to balance differential *versus* equal epigenome establishment in asymmetrically *versus* symmetrically dividing cells, which could then impact plasticity *versus* fidelity in cell fate decisions during development, homeostasis and tissue regeneration.

## Acknowledgments

We thank Chen lab members for insightful suggestions. We thank Isaiah Gao for technical assistance. We thank Johns Hopkins Integrated Imaging Center for confocal imaging. Supported by NIH 5T32GM007231 (J.S., B.D., and J.B.), F31 HD104526 (J.S.), F31 GM115149 (M.W.), NIH R35 GM127075 and R01 HD102474, and the Howard Hughes Medical Institute (X.C.).

## Author Contributions

J.S., B.D., R.R., M.W. and X.C. conceptualized the study. J.S., B.D., R.R., M.W., and J.B. performed all experiments and data analyses. J.S., B.D., R.R., M.W. and X.C. wrote the manuscript. B.D., R.R. and X.C. revised the manuscript.

## Supplementary Materials

Materials and Methods

Figures S1 to S4

Tables S1 to S14

Supplemental References

## Materials and Methods

### Fly strains and husbandry

Fly strains were raised on standard Bloomington media. All flies were raised at 25°C unless noted otherwise. The following fly strains were used: *hs-flp* on the X chromosome (Bloomington Stock Center BL-26902), *nos-Gal4 (with VP16)* on the 2^nd^ chromosome (*1*), *nos-Gal4* (*without VP16* or *DVP16*) on the 2^nd^ chromosome [(from Yukiko Yamashita, Whitehead Institute, USA) and used in (*2*)], *bam-Gal4* on the 3^rd^ chromosome (*3*), *bam-Gal80* on the 3^rd^ chromosome (from Juliette Mathieu and Jean-René Huynh, Collège de France, France), *UASp-FRT-H3-EGFP-FRT-H3-mCherry* on the 2^nd^ chromosome as reported previously (*4*), *polα50* P-element insertion (BL-27205), *polα180* P-element insertion (BL-31805), *pcna>EGFP-pcna* and *rpa>rpa-EGFP* [from Eric Wieschaus, Princeton University, USA and used in (*5*)].

The *polα50* P-element insertion (BL-27205) was verified by sequencing to be a null allele using the following primers: 5’-AGCTCCAATCGTGTATCTCTCT-3’ (specific to the 5’ UTR of the *polα50* gene locus) and 5’-CAATCATATCGCTGTCTCACTC-3’ (specific to the P-element sequences of the EP insertion) were used to amplify the genomic sequences corresponding to the 5’ end of the *polα50* gene locus, where the P-element insertion was located based on the Flybase (https://flybase.org/). Sequencing with this pair of primers confirmed that the P-element is inserted at a position nine base pairs downstream of the start codon, resulting in the new coding sequence 5’-ATGCCCGAAcatgatgaaataaca<u>taa</u> (lowercase sequences indicate the P-element insertion). This leads to eight codons followed by a stop codon (underlined). Hence, this allele results in an early stop codon that very likely represents a null *loss-of-function* allele of the *polα50* gene. This allele is not homozygous viable and is maintained as a heterozygous stock over a balancer chromosome. All experiments using the *polα50^+/-^* background were outcrossing the *polα50*/*Balancer* stock to a *wild-type* stock to have the *polα50* P-element insertion allele over a wild-type chromosome.

### Generating knock-in fly strains

Endogenously tagged fly strains were generated by CRISPR-Cas9 with the genome editing service provided by Fungene Inc. (Beijing, China). The knock-in strains encoding the following proteins were generated and used in this study: Cdc45-mCherry (internally tagged between D163 and Q164), Cdc45-3×HA (internally tagged between D163 and Q164), DNA polymerase ε 255kD subunit-3×HA (tagged at the C-terminus), DNA polymerase α 180kD-3×HA (tagged at the C-terminus), DNA polymerase δ-3×HA (tagged at the C-terminus), Ctf4-EGFP (tagged at the C-terminus).

### Heat shock scheme

Flies with *UASp-FRT-H3-EGFP-FRT-H3-mCherry* along with any relevant genotypes were crossed with *hs-flp; nanos-Gal4* and raised at 25°C. Within two days of eclosure, adult male flies were transferred to a vial and the vial was submerged underwater at 37°C for 90 minutes. Flies were then recovered at 29°C for 18 hours prior to dissection for experiments, with the exception of experiments using the PolA1 inhibitor, as described below.

### Whole mount immunostaining experiments

Immunostaining experiments were performed using standard procedure (*6*). Primary antibodies used were Armadillo (Arm, 1:100; DSHB N2 7A1), Traffic Jam (Tj, 1:100, from Mark Van Doren, Johns Hopkins University, USA), anti-PCNA (1:100; Santa Cruz sc-56), anti-GFP (1:1,000; Abcam ab 13970), anti-HA (1:200; Sigma-Aldrich H3663), anti-mCherry (1:1,000; Invitrogen M11217), anti-H3K27me3 (1:400; Millipore 07-449), anti-H4K20me2/3 (1:400; Abcam ab78517), anti-H3S10ph (1:2000; Cell Signaling Technology 9701), rabbit anti-H3T3ph (1:200, Millipore 05-746R), and anti-BrdU (1:200; Abcam ab6326). BrdU analog was Invitrogen B23151 5-bromo-2′-deoxyuridine (BrdU). Secondary antibodies were the Alexa Fluor-conjugated series (1:1,000; Molecular Probes). Confocal images were taken on the Zeiss LSM800 (with Airyscan mode) with a 63x oil objective lenses or on the Leica SPE with 63x oil immersion lenses.

### Quantification of protein levels in the early germline

Images were analyzed using the ImageJ software FIJI. Germline cyst stages were identified using Arm signal to label the two cyst cells encapsulating each cyst. Average intensity values were recorded for the center Z-slice of each cell/nucleus of interest. For germ cells within one cyst, only one germline nucleus from the entire cyst was measured as one data point. For the comparison of protein levels of endogenously tagged proteins, immunostaining signals in GSCs, 4-cell and 8-cell SGs were measured, and a background was subtracted using the post-mitotic hub cells, which are devoid of signals from any of these replication components. Signal intensity from 4-cell and 8-cell SGs were then normalized to the average intensity of GSCs from the same batch of testes. For the batch-based normalization, within one experimental batch, each data point is normalized to the average of WT GSCs in this corresponding batch. To compare data among different batches, the resulting values were then used to calculate the relative amount of GSC protein level to SG protein level (set to 1 to facilitate comparison), and plot on a log_2_ scale (Fig. 1b). The dataset shown in Figure 1b are from germ cells at each corresponding differentiation stages (Table S1). We also labeled S-phase germ cells using a EdU pulse and quantified them separately. The results using S-phase germ cells were similar to those using germ cells without distinguishing S-phase from G2-phase (data not shown).

For the comparison of the stage-specificity of each driver or driver combination, *nanos-Gal4* by itself, *nos-Gal4ΔVP16; bam-Gal80* combination, or *bam-Gal4* by itself was crossed to the *UASp-FRT-H3-EGFP-FRT-H3-mCherry* transgene without *hs-flp*. The EGFP signals reflecting the relative strength of each driver or driver combination were quantified in the corresponding germline cyst stages, identified using Arm to label the two encapsulating cyst cells. The central slice of a representative nucleus was taken for each cyst measured as one data point. The cytoplasmic space was used as a background for subtraction. The EGFP signals were normalized to the stage with the highest relative signal intensity: For *nanos-Gal4* by itself, all quantifications were normalized to the signals in GSCs; for the *nos-Gal4ΔVP16; bam-Gal80* combination, all quantifications were also normalized to the signals in GSCs; for *bam-Gal4* by itself, all quantifications were normalized to the signals in the 8-cell SGs (Fig. S1e).

### S-Phase colocalization imaging and analysis

To visualize potentially differential histone incorporation during S-phase, we applied a clearance buffer which effectively removes nucleoplasmic protein as previously described (*7, 8*). Briefly, the clearance buffer is prepared by mixing 989µls of the clearance buffer stock solution (8.4 mM HEPES, 100 mM NaCl, 3 mM MgCl, 1 mM EGTA, 300 mM Sucrose, 2% Triton X-1000, and 2% BSA in ddH_2_O) with 1 µl DTT and 10 µl protease inhibitor (100x Leupeptin). After dissection, tissue samples were incubated in 10 μM EdU (Invitrogen Click-iT EdU Imaging Kit, catalog # C10340) for 15 minutes in Schneider’s media at room temperature. At the end of the 15 minutes, the Schneider’s media were drained and the clearance buffer was added for two minutes at 4°C in darkness.

Samples were then fixed in 4% PFA, washed with 1xPBST, and then blocked in 3% BSA for 30 minutes. For robust signals, both the old H3-EGFP and new H3-mCherry were immunostained with antibodies (e.g., anti-EGFP and anti-mCherry) using standard procedures. The CLICK reaction was performed according to manufacturer’s instructions to label EdU. The DNA dye Hoechst was also added at this step.

Images were acquired on the Zeiss LSM800 using Airyscan mode on a 63x oil immersion objective. All samples were imaged using the identical settings. GSCs were identified by their proximity to the hub region. When 4-cell stage SGs were used, only one SG per cyst was analyzed to represent one data point. All images were analyzed using FIJI software. The Pearson score was recorded using the Coloc2 plugin for each nucleus, which was cropped to include just the nucleus as much as possible as delineated by the Hoechst signals. For each batch of images, the average measurement of the control GSCs was set to 1 and the other treatments are normalized to control GSCs, in order to avoid batch variability. The resulting values are then used to calculate mean and standard error of the mean (Mean± SEM).

### Inhibitor treatment and analysis

For S-phase colocalization experiments using the inhibitor, flies were heat shocked as described above and left at 29°C to recover for 14 hours. Testes were then dissected and placed in incubation media for four hours, resulting in 18 total hours of post-heat shock recovery. After incubation with the inhibitor at the designated concentrations, these tissues were processed for S-phase colocalization analysis as described above.

Polα180 inhibitor (MedChemExpress Cat# HY-147812), a derivative of the classical inhibitor adarotene, was prepared in DMSO as stock and stored at −20°C (for short term) and - 80°C (for long term) according to manufacturer’s instructions. Drug incubation was performed on testes in “live cell media” containing Schneider’s insect medium with 200 μg/ml insulin, 15% FBS by volume, and 0.6x pen/strep (*9*). Prior to experiments, incubation media was prepared by diluting inhibitor solution (or DMSO vehicle) to the proper concentration in live cell media. Testes were dissected and placed in 100μl of incubation media as quickly as possible following dissection. Incubated testes were left in open tubes in darkness at room temperature (RT) for four hours. Because four hours are longer than the standard S-phase of the early male germline (*10–14*), all S-phase cells at the end of the incubation should have been exposed to the inhibitor for the entirety of their current S-phase.

For EdU incorporation, 20μM EdU was added to the incubation media for the last 15 minutes of the drug incubation before tissue fixation. Only cells in early-to mid-S-phase were used for quantifications, as denoted by EdU staining covering all or most of the nucleus. Cells with focal EdU signal, indicative of late S-phase, were excluded to avoid skewing of the data. Germ cells were determined by endogenously tagged Vasa-mApple signals. Following imaging, EdU incorporation was quantified by measuring the mean EdU signal intensity in EdU-positive germline nuclei and subtracting the background measured from the nearby EdU-negative cells. When a cyst was considered, only one nucleus from each cyst was measured as one data point. Data shown in Figure S2b were based on all early-stage germ cells, as no significant difference of EdU incorporation was detected among GSCs, GBs, and SGs from the same sample (data not shown).

### Generation of chromatin fibers from the *Drosophila* male germline

Chromatin fibers were prepared as previously described (*4, 15*). Briefly, after adding EdU to the testis samples and incubating for 15 minutes, lysis buffer was added (100 mM NaCl, 25 mM Tris-base, 0.2% Joy detergent, pH=10). The testis tip was then micro-dissected on the slide and the rest of the testis was removed. Cells were allowed to fully lyse for approximately 5 minutes and then a Sucrose/Formalin (1M sucrose; 10% formaldehyde) solution was added and left for 2-minute to incubate, before a cover slip was gently placed on the top. The slide was then transferred to liquid nitrogen for two minutes before the cover slip was removed. The slide was then transferred to 95% EtOH for 10 min at −20°C in a freezer. Afterwards, the slide was fixed in 1% PFA for 1 minute. Samples were washed 3× in a Coplin jar with 1×PBST followed by blocking the sample with 3% BSA in 1×PBST for 30 minutes. Primary antibodies were then added for overnight incubation in a humidity chamber at 4°C. To assess histone asymmetry, anti-PCNA, anti-H3K27me3, and anti-GFP primary antibodies were added to chromatin fibers from the testes from the males with the following genotypes: each of the drivers (*nos-Gal4* itself, *nos-Gal4ΔVP16; bam-Gal80* combination, or *bam-Gal4* itself) crossed with *UASp-FRT-H3-EGFP-FRT-H3-mcherry* without *hs-flp*. For *cdc45-mCherry; DNA Polymerase-HA* fibers, mCherry and HA primary antibodies were used. After the incubation with the primary antibodies, the slides are washed in a coplin jar with 1×PBS. Then the secondary antibodies were added and incubated for two hours at room temperature in a humidity chamber. The click chemistry was performed to label EdU following the manufacturer’s instruction. When DNA needs to be labeled, Hoechst is included at 1:1,000 to stain the samples. Additionally, for samples that need DNA labeling, ProLong™ Gold Antifade Mountant with DNA Stain DAPI (Thermo Fisher catalog # P36931) was used. For samples that do not need DNA labeling, ProLong Diamond mounting media without DAPI (Thermo Fisher catalog# P36961) was used.

### Sequential labeling using EdU and BrdU analogs on DNA fibers

After sample dissection, 10 μM EdU was added for a 10-minute incorporation, followed by washing out EdU. BrdU was subsequently added for another 10 minutes. After this sequential labeling, DNA fibers were prepared using the same procedure as described above for chromatin fibers, with the exception of using a different lysis buffer to strip proteins from the DNA (200 mM Tris–HCl, pH 7.5, 50 mM EDTA, 0.5% SDS). The fibers were then treated with 1M HCl for 30 minutes at room temperature to expose the incorporated BrdU. After washing with 1×PBST, BrdU antibodies were added for incubation overnight at 4°C in a humidity chamber. Secondaries antibodies against the BrdU primary antibodies were then added for two hours at room temperature in a humidity chamber. The click reaction to recognize EdU was performed subsequently along with Hoechst incubation at 1:1000. Samples were then mounted in ProLong Diamond mounting media with DAPI. The EdU-positive DNA fibers representing regions that undergo DNA replication during EdU pulse (and thus have EdU on at least one side) were used for subsequent analyses as shown in Figure 5b.

### Identifying and imaging replicative DNA fibers and chromatin fibers

All DNA fibers and chromatin fibers in this study were imaged with the Airyscan mode on a Zeiss LSM800 using a 63× oil immersion lens. Germline-derived chromatin fibers were identified using the H3-EGFP signal expressed with different germ cell-specific drivers or driver combination. Replicative regions were identified by both PCNA and EdU signals, or the presence of Cdc45, DNA Polymerase, and EdU. Fibers regions with detectable separation between sister chromatids were imaged and analyzed. Quality controls to select appropriate chromatin fiber regions for further analyses included fiber length, shape, and the molecular specificity of signals. For example, for quantifying old histone-enriched H3K27me3 with strandedness information, the EdU labeled fibers positive with PCNA, H3-EGFP and H3K27me3 signals were used. For analyzing the Cdc45 signals with DNA polymerases, fibers with EdU-labeling regions, clear Cdc45 and anti-HA signals were used.

For sequential EdU and BrdU labeled DNA fibers, two patterns were imaged and quantified at DNA regions that replicate during the EdU pulse (thus incorporating EdU on at least one side of the duplicated sister chromatids): First, regions with clear sister chromatid separation with Hoechst and EdU signals but no discernable BrdU signal. Second, regions with clear sister chromatid separation with clear Hoechst, EdU, and BrdU signals. For detailed description of the analyses of sister chromatids using chromatin fibers, refer to (*4, 15*).

### Quantification of DNA fibers and chromatin fibers

All images were analyzed using FIJI software. To quantify the asymmetry between sister chromatids, line plots were drawn on both strands, using the PCNA-enriched side to denote the lagging strand. Most fibers have relatively short separable regions (≤ 2μm), for which the entire fiber was used for quantification. For fibers with longer separable regions (> 2 μm), they were divided into 2μm-long non-overlapping segments along the length of the chromatin fiber and each of them was used for analyses. The region with no overlap with any of the chromatin fibers was used as background signal for subtraction from the measured signals from both strands. The ratio of signals = log_2_ (leading strand signal ̶ background signal) / (lagging strand signal ̶ background signal).

For the sequential EdU- and BrdU-labeled DNA fibers, there is no strandedness indicator such as PCNA. As such, the strand with higher BrdU signals was used as the reference strand, allowing EdU signal to be independently measured, which could be on the same or the opposite strand. All quantifications were performed similar to the chromatin fibers, with the ratio of signals = log_2_ (BrdU-enriched strand signal ̶ background signal) / (BrdU-depleted strand signal ̶ background signal).

For the Cdc45- and DNA Polymerase-labeled fibers, the distance between Cdc45 signal and the HA signal (labeling either Polα or Polε) was quantified from the center of the Cdc45 focus to the nearest HA signal.

### A quantitative assay for chromosomal condensation state

We used an area-based method to monitor the chromosomal condensation state as previously described (*8*), using a dual-color histone transgene *UASp-FRT-histone-EGFP -FRT-histone-mCherry*. A maximum intensity projection was generated for old H3- (EGFP) and new H3- (mCherry) enriched areas. The intensity of each pixel was determined and scaled individually, setting the minimum intensity to 0 and the maximum to 65,535 (a 16-bit range). We monitored the pixels across the image with a threshold of 35% of the maximum intensity. Condensation kinetic profiles were generated to compare old H3- *versus* new H3-enriched regions by calculating the percentage of pixels above the threshold (the condensation parameter). Relative compaction index was measured and plotted by taking a ratio of the percentage of pixels of the new H3-enriched to the old H3-enriched regions as described previously (*8*).

### Statistics and reproducibility

For all comparisons between two groups, Mann-Whitney tests were used unless otherwise noted. For one-group datasets, one sample *t*-test was used with a null hypothesis that the data is symmetrically distributed (e.g., ratio= 1 for datasets without logarithmic transformation, log_2_= 0 for logarithmically transformed data).

### Details for **Figure 1b**

Endogenously expressed Polα-HA levels are significantly depleted in GSCs relative to SGs according to the Mann-Whitney test with a *P*-value < 10^-4^ (****).

Endogenously expressed Polδ-HA levels are significantly depleted in GSCs relative to SGs according to the Mann-Whitney test with a *P*-value < 10^-4^ (****).

Endogenously expressed Polε-HA levels are not significantly different between GSCs and SGs according to the Mann-Whitney test with a *P*-value > 0.05 (= 0.0806, ns).

Endogenously expressed Cdc45-HA levels are not significantly different between GSCs and SGs according to the Mann-Whitney test with a *P*-value > 0.05 (= 0.8469, ns).

Endogenously expressed Ctf4-EGFP levels are not significantly different between GSCs and SGs according to the Mann-Whitney test with a *P*-value > 0.05 (= 0.8595, ns).

RPA driven under its own promoter (*rpa>rpa-EGFP*) levels are significantly enriched in GSCs relative to SGs according to the Mann-Whitney test with a *P*-value < 10^-4^ (****).

### Details for Fig. 3f and Fig. S3c

*nos-Gal4ΔVP16; bam-Gal80>H3-EGFP* (GSC-enriched) chromatin fibers exhibited significantly higher levels of H3K27me3 asymmetry relative to *nanos-Gal4>H3-EGFP* (total early germline) chromatin fibers according to the Mann-Whitney test with a *P*-value < 0.05 (= 0.0250, *).

*bam-Gal4>H3-EGFP* (SG) chromatin fibers exhibited significantly lower levels of H3K27me3 asymmetry relative to *nanos-Gal4>H3-EGFP* (total early germline) chromatin fibers according to the Mann-Whitney test with a *P*-value < 0.05 (= 0.0296, *).

The *in silico* combination of *nos-Gal4ΔVP16; bam-Gal80>H3-EGFP* (GSC-enriched) and *bam-Gal4>H3-EGFP* (SG) chromatin fibers was not statistically different from the *nanos-Gal4>H3-EGFP* (total early germline) chromatin fibers according to the Mann-Whitney test with a *P*-value > 0.05 (= 0.7458, ns).

*nos-Gal4>H3-EGFP*; *polα50^+/-^* chromatin fibers exhibited significantly higher levels of H3K27me3 asymmetry relative to *nanos-Gal4>H3-EGFP* chromatin fibers according to the Mann-Whitney test with a *P*-value < 0.05 (= 0.0256, *).

*bam-Gal4>H3-EGFP*; *polα50^+/-^* chromatin fibers exhibited significantly higher levels of H3K27me3 asymmetry relative to *bam-Gal4>H3-EGFP* chromatin fibers according to the Mann-Whitney test with a *P*-value < 0.01 (= 0.0075, **).

*nos-Gal4ΔVP16; bam-Gal80>H3-EGFP*; *polα50^+/-^*chromatin fibers were not statistically different from the *nos-Gal4ΔVP16; bam-Gal80>H3-EGFP* chromatin fibers according to the Mann-Whitney test with a *P*-value > 0.05 (= 0.8039, ns).

### Details for Fig. S3d

*nos-Gal4>H3-EGFP*; *polα180^+/-^* chromatin fibers exhibited significantly higher levels of H3K27me3 asymmetry relative to *nanos-Gal4>H3-EGFP* chromatin fibers according to the Mann-Whitney test with a *P*-value < 0.01 (= 0.0047, **).

*nos-Gal4>H3-EGFP; rpa70-HA* chromatin fibers exhibited significantly higher levels of H3K27me3 asymmetry relative to *nanos-Gal4>H3-EGFP* chromatin fibers according to the Mann-Whitney test with a *P*-value < 0.05 (= 0.0209, *).

### Details for Fig. S4b

For DNA fibers with both EdU and BrdU signals, EdU and BrdU exhibit significantly different distribution according to the Mann-Whitney test with a *P*-value < 10^-4^ (****).

For DNA fibers with both EdU and BrdU signals, BrdU is significantly asymmetrically localized by a one sample t-test with a null hypothesis of log_2_=0 and a *P*-value < 10^-4^ (****).

For DNA fibers with both EdU and BrdU signals, EdU is significantly asymmetrically localized by a one sample t-test with a null hypothesis of log_2_=0 and a *P*-value < 10^-4^ (****).

### Details for Figure 5g

The chromatin fibers co-labeled with Cdc45-mCherry and Polα-HA exhibit significantly longer distances between mCherry focus and HA signal relative to the chromatin fibers co-labeled with Cdc45-mCherry and Polε-HA by the Mann-Whitney test with a *P*-value < 10^-4^ (****).

### Details for Fig. S4d

In *nos-Gal4>H3-EGFP* chromatin fibers, H3K27me3 is significantly asymmetrically localized by a one sample t-test with a null hypothesis of log_2_= 0 and a *P*-value < 10^-4^ (****).

In *nos-Gal4>H3-EGFP* chromatin fibers, PCNA was significantly asymmetrically localized by a one sample t-test with a null hypothesis of log_2_= 0 and a *P*-value < 10^-4^ (****).

In *nos-Gal4>H3-EGFP* chromatin fibers, EdU was not significantly asymmetrically localized by a one sample t-test with a null hypothesis of log_2_= 0 and a *P*-value > 0.05 (= 0.1028, ns).

## Supplemental Figures and Figure Legends

**Figure S1:**
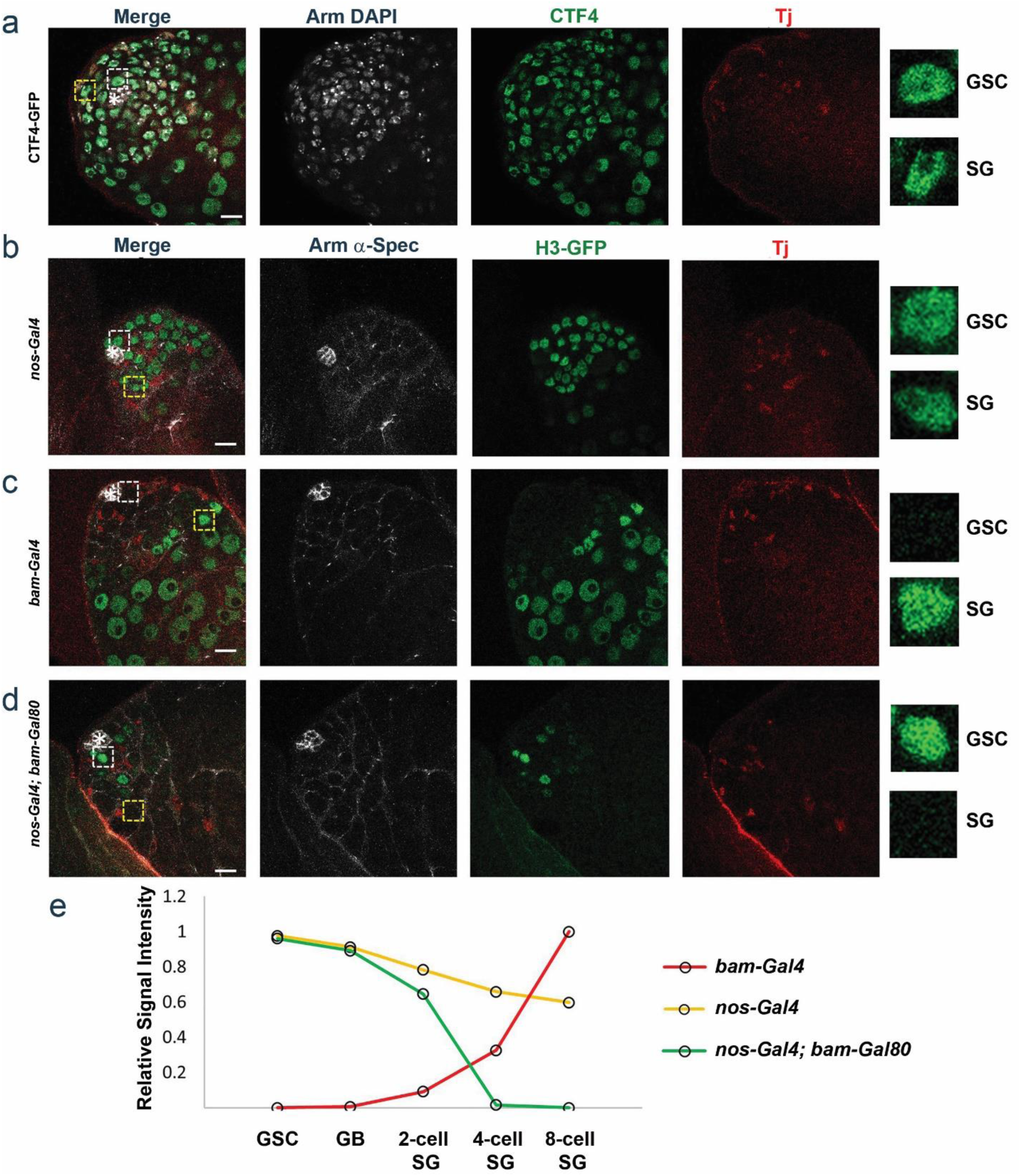
Expression pattern of a replication machinery component CTF4 and distinct expression patterns of an H3-EGFP reporter by different drivers in the *Drosophila* male germline. (**a**) Image of endogenous CTF4-GFP using knock-in strategy (Materials and Methods): DAPI (white), Arm (white, a marker for hub cells), CTF4-GFP (green), and the somatic marker Traffic Jam (Tj, red). (**b-d**) Images of: (**b**) *nanos-Gal4> UAS-H3-EGFP*, (**c**) *bam-Gal4> UAS-H3-EGFP*, and (**d**) *nos-Gal4ΔVP16; bam-Gal80> UAS-H3-EGFP*, Arm (white), H3-EGFP (green), and the somatic marker Tj (red). (**e**) Quantification of the relative expression levels of H3-EGFP using each corresponding driver (n=3 for each genotyped testes, see Table S2 for details). Asterisk: hub. Scale bar: 10 μm.

**Figure S2:**
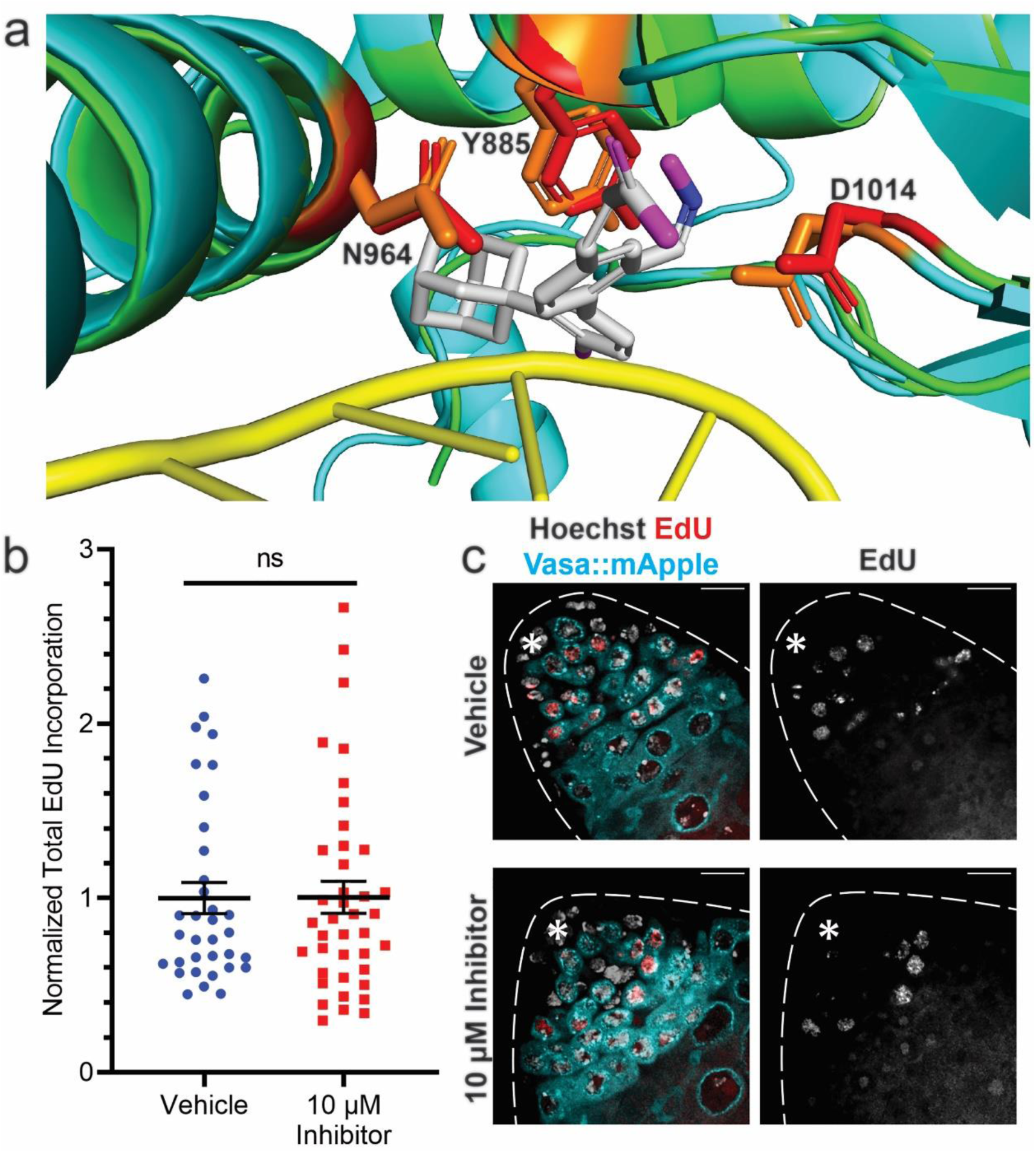
Low concentration of Polα180 (or PolA1) inhibitor partially inhibits Polα while permitting DNA replication. (**a**) PyMOL structural alignment of human DNA Polα (green) with DNA (yellow) (structure: PDB 5IUD) and the AlphaFold prediction of *Drosophila* Polα180 (cyan). The Polα180 inhibitor (carbon: white, oxygen: magenta, nitrogen: blue) is shown at the binding location predicted by (*16*). The Polα180 residues predicted to interact with the inhibitor are conserved between mammals (red) and *Drosophila* (orange). (**b**) Quantification of total EdU incorporation in early-stage germ cells, treated with vehicle or Polα180 inhibitor for four hours, normalized to the mean of vehicle-treated cells. Vehicle treated cells (n=34), 10μM inhibitor treated cells (n=40). Median with first and third quartile shown. Student’s t-test, ns: not significant. See Table S5 for details. (**c**) Representative images of testes treated with vehicle or Polα180 inhibitor. In merged images: Hoechst (white), endogenous Vasa-mApple (cyan), EdU (red). Asterisk: hub. Scale bars: 10 μm.

**Figure S3:**
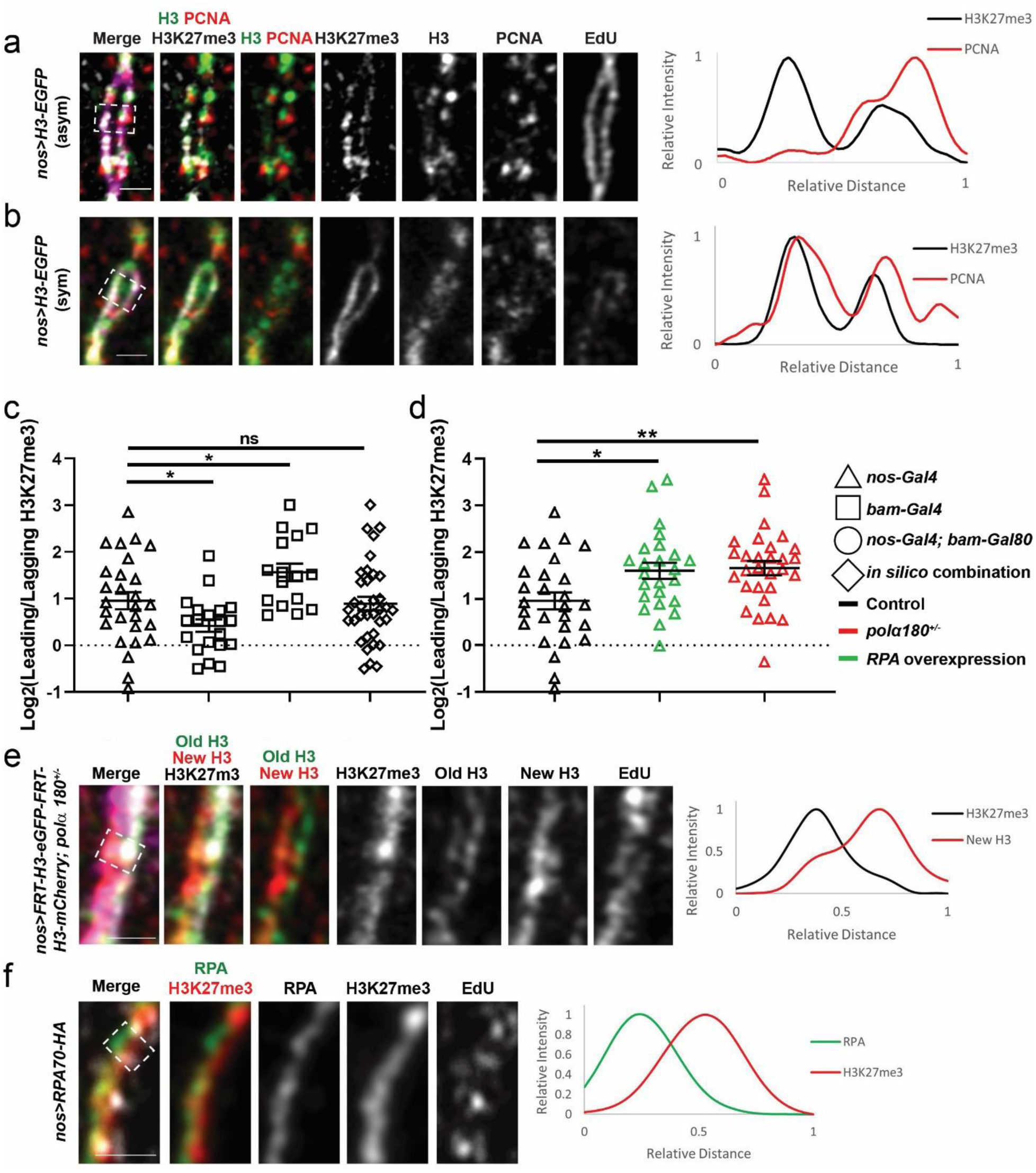
Reducing Polα or enhancing RPA levels increase old histone-enriched H3K27me3 asymmetries at the replication fork. (**a**) An Airyscan image of representative asymmetric *nanos-Gal4>H3-EGFP* chromatin fiber. (**b**) An Airyscan image of representative symmetric *nanos-Gal4>H3-EGFP* chromatin fiber. In merged images (**a**-**b**): H3K27me3 (white), H3-EGFP (green), PCNA (red), and EdU (magenta). Images in (**a**-**b**) are also accompanied by line plots showing the spatial distribution of H3K27me3 and PCNA signals from the indicated regions, respectively (white dotted outlined box). (**c**) Quantification of H3K27me3 asymmetry using chromatin fibers with *nanos-Gal4>H3-EGFP*, *bam-Gal4>H3-EGFP*, *nos-Gal4ΔVP16*; *bam-Gal80>H3-EGFP*, and an *in silico* combination of *bam-Gal4>H3-EGFP* and *nos-Gal4ΔVP16*; *bam-Gal80>H3-EGFP* in log_2_ scale: *nanos-Gal4>H3-EGFP*= 0.95± 0.18 (n=27), *bam-Gal4>H3-EGFP*= 0.42± 0.14 (n=20), *nos-Gal4ΔVP16*; *bam-Gal80>H3-EGFP*= 1.56± 0.19 (n=16), *in silico* combination of *bam-Gal4>H3-EGFP* and *nos-Gal4ΔVP16*; *bam-Gal80>H3-EGFP*= 0.89± 0.15 (n=36). See Table S6 for details. (**d**) Quantification of additional replication protein manipulations using a *nanos-Gal4* driven overexpression of *UAS-rpa70-HA* transgene and a P-element insertion allele of another Polα subunit gene (*polα180*) at a heterozygous background (*polα180^+/-^*) in log_2_ scale: *nanos-Gal4>rpa70-HA*= 1.60± 0.17 (n=24), *nanos-Gal4>H3-EGFP; polα180^+/-^*= 1.66± 0.15 (n=29). See Table S7 for details. (**e**) Airyscan image of *hs-flp*; *nanos-Gal4*>*FRT-H3-EGFP-FRT-H3-mCherry*; *polα180^+/-^*: H3K27me3 (white), old H3 (green), new H3 (red), and EdU (magenta) in the merged image. (**f**) Airyscan image of *nanos-Gal4>rpa70-HA* chromatin fiber: EdU (white), RPA (green), and H3K27me3 (red) in the merged image. Images in (**e**-**f**) are also accompanied by line plots showing the spatial distribution of H3K27me3 and new H3 signals in (**e**), as well as H3K27me3 and RPA signals in (**f**) from the indicated regions, respectively (white dotted outlined box). Scale bar: 1 μm. All ratios: Mean± SEM. All statistics: Mann-Whitney test, **: *P*< 0.01, *: *P*< 0.05, ns: not significant.

**Figure S4:**
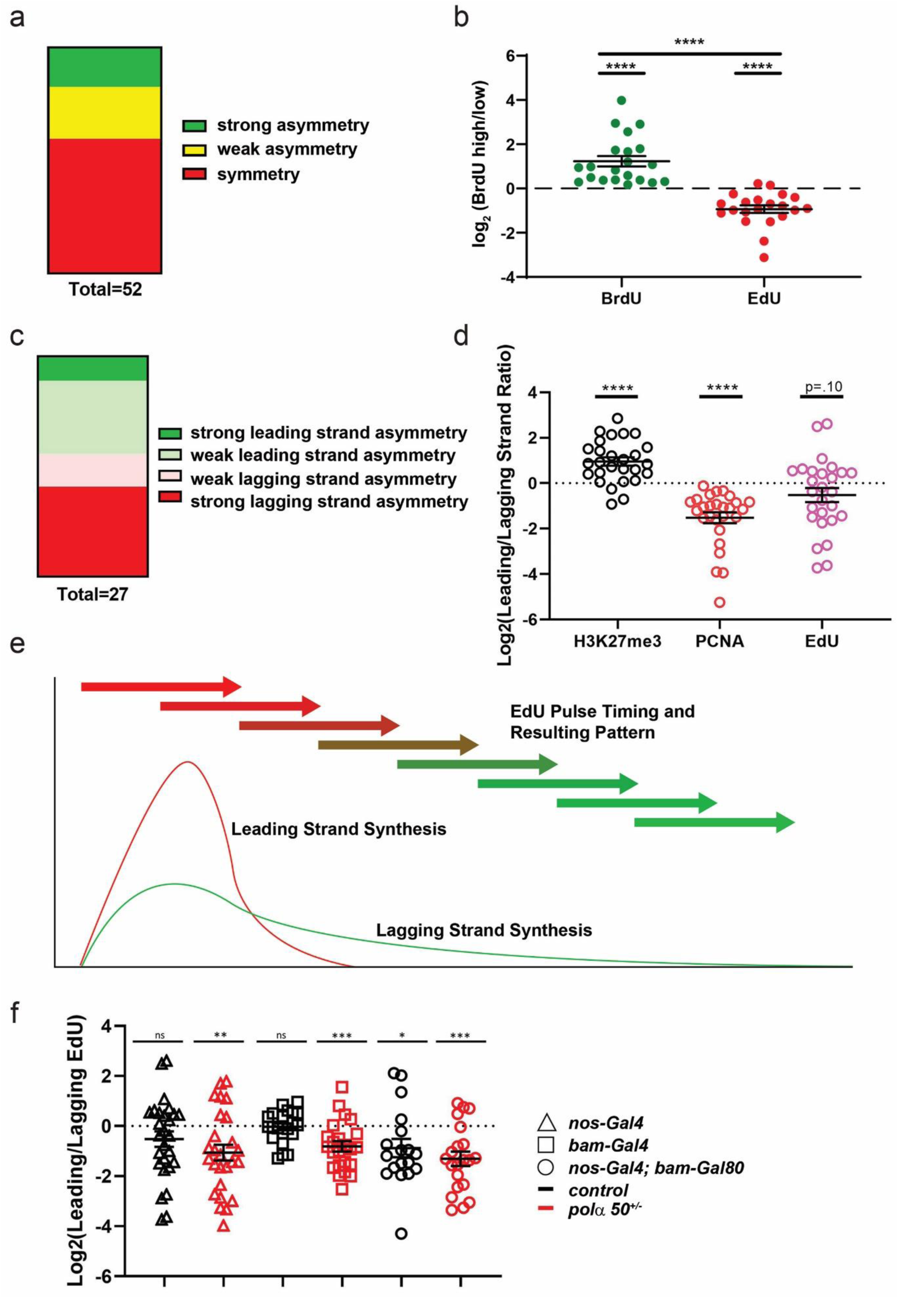
Visualization and quantification of delayed lagging strand synthesis. (**a**) Assessment of observed patterns on DNA fibers, wherein symmetric fibers refer to fibers with EdU on both strands, weak asymmetric fibers have both EdU and BrdU but less than a 2-fold asymmetry on both strands, while strong asymmetric fibers have a greater than 2-fold difference for at least one of the signals (i.e., either EdU or BrdU). The distribution of the three categories are as follows: 60% symmetric, 23% weak asymmetric, 17% strong asymmetric. See Table S9 for details. (**b**) Log_2_-scale 1D quantification of EdU and BrdU from the DNA fibers with both signals, where the positive side is the strand with higher BrdU and the negative side is the strand with higher EdU: log_2_BrdU= 1.23± 0.23 (n=21), log_2_EdU= −0.93± 0.17 (n= 21). ****: *P*< 10^-4^, Mann-Whitney test for the comparison between two groups, one tailed t-test with a null hypothesis of log_2_= 0 (symmetric pattern). See Table S10 for details. (**c**) Assessment of EdU asymmetries wherein ≥ 2-fold are considered strong asymmetry, < 2-fold are considered weak asymmetry. The distribution of the three categories are as follows: 11% strong asymmetry toward the leading strand, 33% weak asymmetry toward the leading strand, 15% weak asymmetry toward the lagging strand, 41% strong asymmetry toward the lagging strand. See Table S13 for details. (**d**) Quantification of H3K27me3, PCNA, and EdU asymmetry from *nos-Gal4>H3-EGFP* labeled chromatin fibers using log_2_ scale: log_2_H3K27me3= 0.95± 0.18 (n=27), log_2_PCNA= −1.52± 0.24 (n=27), log_2_EdU= −0.52± 0.31 (n=27). ****: *P*< 10^-4^, one tailed t-test with a null hypothesis of log_2_= 0 (symmetric pattern). See Table S14 for details. (**e**) A model of replication patterns that could explain the observed EdU patterns. (**f**) Quantification of EdU distribution on chromatin fibers labeled with H3-EGFP driven by the following drivers with regard to the strandedness using log_2_ scale: *nanos-Gal4*= −0.52± 0.31 (n=27, *P*= 0.103, ns), *nanos-Gal4*; *polα50^+/-^*= −1.06± 0.32 (n=26, *P*< 0.01), *bam-Gal4*= −0.03± 0.14 (n=21, *P*= 0.817, ns), *bam-Gal4*; *polα50^+/-^*= −0.081± 0.21 (n=23, *P*< 10^-3^), *nos-Gal4ΔVP16*; *bam-Gal80*= −0.88±0.37 (n=18, *P*< 0.05), *nos-Gal4ΔVP16*; *bam-Gal80*; *polα50^+/-^*= −1.31± 0.29 (n=21, *P*< 10^-3^). All ratios: Mean± SEM, one tailed t-test with a null hypothesis of log_2_= 0 (symmetric pattern), ***: *P*< 10^-3^, **: *P*< 0.01, *: *P*< 0.05, ns: not significant. See Table S11 for details.

## Supplemental Tables

**Table S1:** Raw Data Related to Figure 1b (log_2_ scale):

| GSC Polα | SG Polα | GSC Polδ | SG Polδ | GSC Polε | SG Polε | GSC Cdc45 | SG Cdc45 |
| --- | --- | --- | --- | --- | --- | --- | --- |
| -1.31257 | 0.478214 | -0.53529 | 0.025996 | -0.05733 | 0 | -0.40234 | -0.18844 |
| 0.309191 | 0.179521 | -0.22863 | 0.522421 | 0.108252 | -0.28951 | 0.118644 | -0.34792 |
| -0.37796 | 0.34155 | -0.82902 | -0.28951 | 0.192645 | -0.11704 | -0.18844 | -0.10692 |
| -0.58645 | -0.30091 | -1.13936 | -1.02647 | 0.036995 | 0.303069 | 0.166649 | 0.014646 |
| 0.282692 | -0.70322 | -1.25966 | 0.368388 | -0.1375 | 0 | 0.111576 | -0.02975 |
| -0.8576 | -1.69149 | -1.08278 | 0.172836 | -0.33605 | 0.37707 | -0.25716 | -0.18844 |
| -1.74014 | 0.843809 | -1.02833 | 0.115881 | -0.4339 | 0.272372 | 0.014646 | 0 |
| -1.29675 | -0.41973 | -1.78322 | -0.35509 | 0.125531 | -0.2007 | 0.032029 | -0.00295 |
| -1.39827 | -1.2292 | -1.78322 | -0.73696 | -0.01886 | -0.3599 | 0.166649 | -0.32931 |
| -1.09303 | 0.909612 | -1.46129 | -0.1375 | -0.22239 | -0.48543 | -0.50619 | 0.179706 |
| -2.0808 | 0.249741 | -0.35695 | 0 | -0.56635 | -0.48543 | 0.192645 | 0.099536 |
| -2.41359 | -1.2475 | -0.61329 | 0.466568 | 0.159479 | -0.38414 | 0.057715 | 0.20547 |
| -2.87359 | -0.29352 | -1.19826 | 0 | 0.347923 | -0.28951 | 0.192645 | 0.179706 |
| -1.72364 | -0.34207 | -0.97586 | -0.1375 | 0.036995 | -0.03797 | -0.31093 | -0.0148 |
| -0.10579 | -0.23503 | -0.82902 | -0.2115 | -0.24442 | -0.4339 | -0.20532 | 0.304153 |
| -0.16098 | -0.27503 | -1.08278 | -0.49596 | -0.4088 | -0.5119 | 0.099536 | -0.32931 |
| -1.39736 | 0.492338 | -0.46129 | -0.06711 | -0.46667 | -0.65208 | -0.50773 | 0.166649 |
| -0.9895 | 0.601421 |  | -0.67283 | -0.42182 | -0.31259 | 0.166649 | 0.140178 |
| -0.65638 | 0.397087 |  | -0.08927 | -0.5771 | 0.036995 | 0.071791 | 0.126757 |
| -0.59786 | -0.77119 |  | -0.65703 | -0.73275 | -0.26679 | 0.113211 | 0.065614 |
| -1.33687 | 0.605119 |  | -0.78587 |  | -0.05733 | -0.32931 | -0.44479 |
| -0.58599 | 0.530551 |  | -0.2863 |  | 0.192645 |  |  |
| -0.87321 | 0.278698 |  | -0.08092 |  | 0.287803 |  |  |
| -1.41299 | -1.55611 |  | 0 |  | 0.318177 |  |  |
| -0.55373 | 0.58095 |  | 0.172836 |  | 0.256775 |  |  |
| -0.8475 | -0.91912 |  | 0.125532 |  | 0.303069 |  |  |
| -1.17455 | 0.525471 |  | 0.076621 |  | 0 |  |  |
| -0.84107 | 0.469582 |  | 0.327165 |  | -0.17932 |  |  |
| -0.06675 | 0.312027 |  | 0.149378 |  | 0.272372 |  |  |
| 0.17612 | 0.67072 |  | -0.10893 |  | -0.03797 |  |  |
| 0.604322 | 1.002294 |  | 0.327165 |  | -0.24442 |  |  |
| -0.64542 | 0.194597 |  | 0.558491 |  | -0.1375 |  |  |
| -0.10228 | -0.09784 |  | 0.172836 |  | -0.45943 |  |  |
| -0.49791 | 0.424877 |  | -0.1375 |  | -0.1375 |  |  |
| -2.1112 | 0.425666 |  | 0.742202 |  | 0.36257 |  |  |
| -1.29082 | 0.549028 |  | 0.862497 |  | 0.241008 |  |  |
| -2.74306 | -0.42891 |  | 0.149378 |  | 0.125531 |  |  |
|  | -0.68767 |  | 0.149378 |  | 0.192645 |  |  |
|  | -1.09098 |  | 0.241008 |  | 0.287803 |  |  |
|  | 0.16498 |  | -0.05344 |  | 0.347923 |  |  |
|  | -0.26264 |  | -0.1964 |  | 0.256775 |  |  |
|  | -0.3827 |  | 0.7578 |  | 0.018616 |  |  |
|  | -0.2624 |  | 0.77323 |  | 0.287803 |  |  |
|  | 0.338635 |  | 0.803603 |  |  |  |  |
|  | 0.196455 |  | 0.742202 |  |  |  |  |
|  | 0.017312 |  | 0.803603 |  |  |  |  |
|  | -0.28549 |  | 0.59368 |  |  |  |  |
|  | -1.18792 |  | 0.833351 |  |  |  |  |
|  | 0.486708 |  | 0.77323 |  |  |  |  |
|  | 0.583987 |  | 0.263035 |  |  |  |  |
|  | -0.38879 |  | -1.45943 |  |  |  |  |
|  | -0.38339 |  | -1.38904 |  |  |  |  |
|  | 0.96703 |  | -1.02647 |  |  |  |  |
|  | -1.51239 |  | -1.32193 |  |  |  |  |
|  | -0.35785 |  | -0.65208 |  |  |  |  |
|  | 0.16457 |  | -0.61143 |  |  |  |  |
|  | -0.42837 |  | -1.08092 |  |  |  |  |
|  | -0.91937 |  | -0.92338 |  |  |  |  |
|  | -0.16938 |  | -0.53343 |  |  |  |  |
|  | -0.24721 |  | -1.45223 |  |  |  |  |
|  | -0.32259 |  | 0.412413 |  |  |  |  |
|  | -2.00388 |  | 0.847997 |  |  |  |  |
|  | -0.63923 |  | 0.347924 |  |  |  |  |
|  | -0.93225 |  | 0.051531 |  |  |  |  |
|  | -2.37993 |  | -0.53343 |  |  |  |  |
|  | -0.04774 |  | -0.10893 |  |  |  |  |
|  | -1.16026 |  | -0.6939 |  |  |  |  |
|  | -0.20723 |  | -0.78136 |  |  |  |  |
|  | -1.70132 |  | 0.788496 |  |  |  |  |
|  | -0.61486 |  | 0.803603 |  |  |  |  |
|  | -0.65722 |  |  |  |  |  |  |
|  | 0.211628 |  |  |  |  |  |  |
|  | 0.056212 |  |  |  |  |  |  |
|  | -2.66654 |  |  |  |  |  |  |
|  | -0.89585 |  |  |  |  |  |  |
|  | -1.06545 |  |  |  |  |  |  |
|  | -0.15041 |  |  |  |  |  |  |
|  | -0.32111 |  |  |  |  |  |  |
|  | 0.264392 |  |  |  |  |  |  |
|  | -0.59622 |  |  |  |  |  |  |
|  | 0.09556 |  |  |  |  |  |  |
|  | 0.560539 |  |  |  |  |  |  |
|  | 0.63656 |  |  |  |  |  |  |
|  | 0.314024 |  |  |  |  |  |  |
|  | -0.00704 |  |  |  |  |  |  |
|  | -2.17927 |  |  |  |  |  |  |
|  | -1.23407 |  |  |  |  |  |  |
|  | -0.51216 |  |  |  |  |  |  |
|  | -0.34315 |  |  |  |  |  |  |
|  | -0.6354 |  |  |  |  |  |  |
|  | -0.18619 |  |  |  |  |  |  |
|  | 0.56031 |  |  |  |  |  |  |
|  | 0.374148 |  |  |  |  |  |  |
|  | 0.559079 |  |  |  |  |  |  |
|  | 0.300244 |  |  |  |  |  |  |
|  | -0.49019 |  |  |  |  |  |  |
|  | -1.13866 |  |  |  |  |  |  |
|  | 0.79303 |  |  |  |  |  |  |
|  | 0.440277 |  |  |  |  |  |  |
|  | 0.96703 |  |  |  |  |  |  |
|  | 0.741985 |  |  |  |  |  |  |
|  | -1.01161 |  |  |  |  |  |  |
|  | -0.22325 |  |  |  |  |  |  |
|  | -0.27899 |  |  |  |  |  |  |
|  | 0.875468 |  |  |  |  |  |  |
|  | 0.510143 |  |  |  |  |  |  |
|  | 0.5178 |  |  |  |  |  |  |
|  | 0.713183 |  |  |  |  |  |  |
|  | 0.633194 |  |  |  |  |  |  |
|  | 0.886434 |  |  |  |  |  |  |
|  | 0.439395 |  |  |  |  |  |  |
|  | 0.835981 |  |  |  |  |  |  |
|  | 0.491163 |  |  |  |  |  |  |
|  | 0.432999 |  |  |  |  |  |  |
|  | 0.097461 |  |  |  |  |  |  |
|  | 0.06878 |  |  |  |  |  |  |
|  | 0.095233 |  |  |  |  |  |  |
|  | 0.007007 |  |  |  |  |  |  |

| GSC Ctf4 | SG Ctf4 | GSC RPA | SG RPA |
| --- | --- | --- | --- |
| 0.623142 | 0.951972 | 0.71413 | 0.46296 |
| 0.609015 | -0.50849 | 0.148959 | 0.63969 |
| 0.171094 | -0.1383 | 0.511862 | -0.05705 |
| -0.35686 | -0.74374 | 0.6915 | 0.228219 |
| 0.066299 | 0.490909 | 0.75115 | -0.01768 |
| -0.04728 | -0.26161 | 0.508111 | 0.396194 |
| 0.243402 | 0.802319 | 0.644909 | -0.98263 |
| 0.353389 | -0.05505 | 0.410073 | -0.63155 |
| 0.204584 | 0.254205 | 0.951807 | -0.71856 |
| 0.573534 | -0.79276 | 0.699907 | -0.57468 |
| -0.00298 | 0.196288 | 0.580045 | -0.30973 |
| 0.053226 | 0.364334 | 0.606322 | -2.00897 |
| 0.127204 | -0.37851 | 0.762479 | -2.80474 |
| 0.210356 | 0.278883 | 0.529572 | -1.82867 |
| 0.003056 | 0.452054 | 0.494567 | -0.90634 |
| -0.30331 | -0.29025 | 0.683544 | -0.8102 |
| -0.08452 | -0.07451 | 0.426963 | -0.58593 |
| 0.275981 | 0.105654 | 1.051248 | 0.162835 |
| 0.103751 | 0.704144 | 0.686968 | 0.149163 |
| -0.18246 | -0.3556 | -0.21195 | -0.01045 |
| 0.090752 | -0.16395 | 0.174263 | 0.939597 |
| -0.35899 | 0.116547 | 0.451457 | -0.16126 |
| -0.01312 | 0.362708 | 0.555232 | 0.094393 |
| 0.064503 | 0.040772 |  | 0.021406 |
| -0.1226 | -0.06534 |  | -0.68737 |
| -0.39215 | -0.35847 |  | -0.74938 |
| 0.262724 | 0.742378 |  | 0.174443 |
| -0.01062 | -0.31665 |  | 0.088719 |
| 0.123721 | 0.370067 |  | -0.58799 |
| -0.19653 | -0.4246 |  | -0.45357 |
| 0.134335 | -0.49635 |  | -0.10598 |
| -0.07488 | 0.011934 |  | -0.28448 |
| 0.118006 | -0.29622 |  | 0.457047 |
| 0.68878 | 0.201033 |  | 0.438428 |
| -0.36554 | 0.608532 |  | 0 |
| -0.24475 | -0.42571 |  | 0.239108 |
| 0.013141 | 0.241597 |  | 0.511733 |
| -0.44443 | 0.268592 |  | 0.605868 |
| -0.32045 | -1.04457 |  | 0.445787 |
| -0.27927 | -0.01203 |  | 0.268457 |
|  |  |  | 0.261867 |
|  |  |  | -0.94206 |
|  |  |  | 0.465529 |
|  |  |  | -1.45961 |
|  |  |  | -0.445 |
|  |  |  | -1.92342 |
|  |  |  | -0.26846 |
|  |  |  | -0.78815 |
|  |  |  | -0.07116 |
|  |  |  | 0.389252 |
|  |  |  | 0.200518 |
|  |  |  | 0.615396 |
|  |  |  | 0.309905 |
|  |  |  | 0.642078 |
|  |  |  | 0.611062 |
|  |  |  | 0.38686 |
|  |  |  | 0.141967 |

**Table S2:**
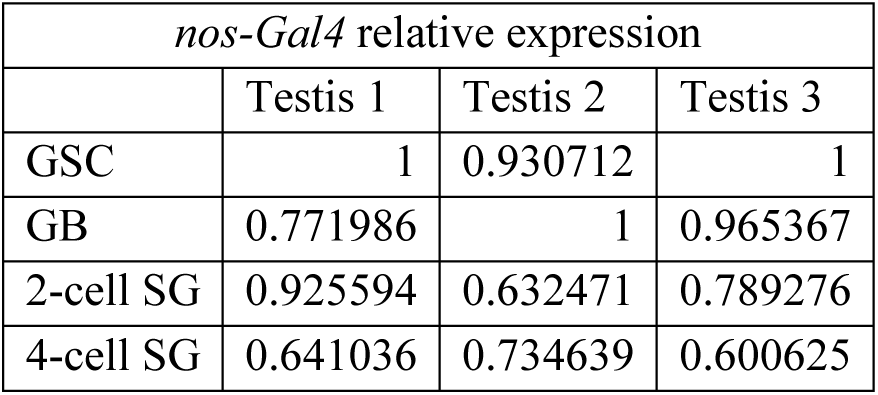
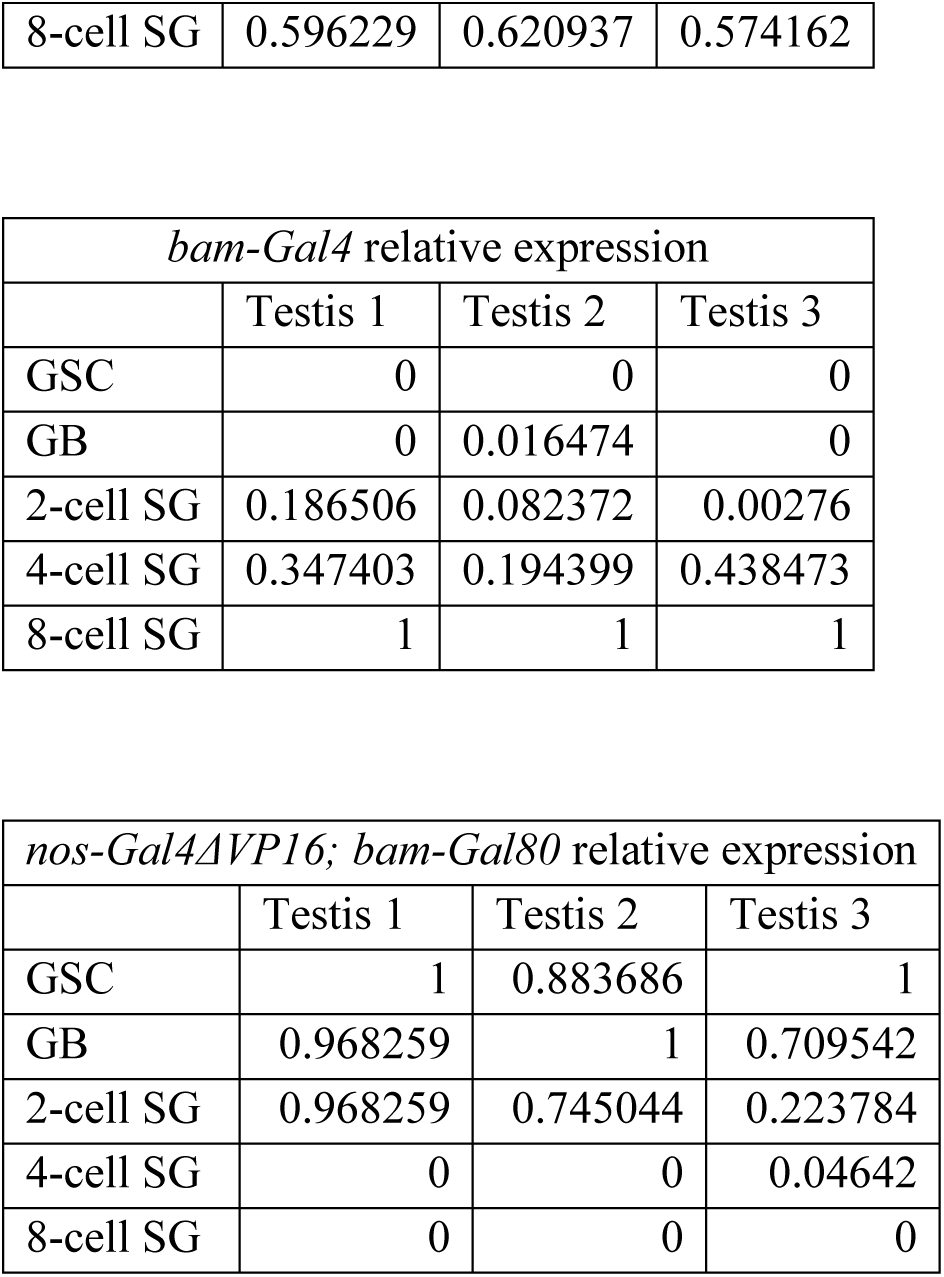
Raw Data Related to Figure S1e:

**Table S3:**
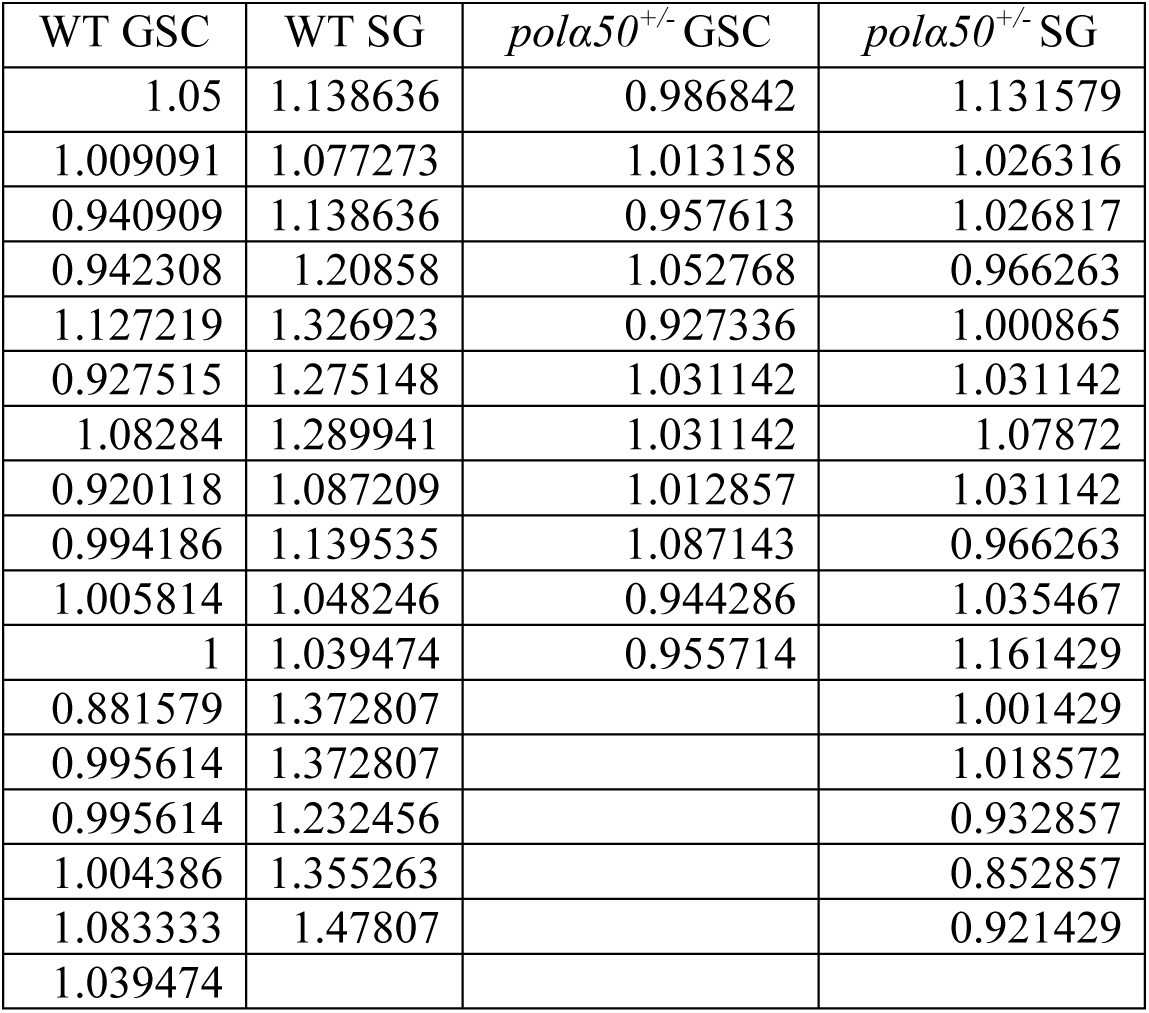
Raw Data Related to Figure 2c:

**Table S4:** Raw Data Related to Figure 2f:

| GSC |  |  |  | SG |  |  |  |
| --- | --- | --- | --- | --- | --- | --- | --- |
| Vehicle | 2.5µM inhibitor | 5.0µM inhibitor | 10µM inhibitor | Vehicle | 2.5µM inhibitor | 5.0µM inhibitor | 10µM inhibitor |
| 1.109244 | 1.092308 | 1.051546 | 1.086792 | 1.092437 | 0.969231 | 0.943299 | 1.207547 |
| 0.890756 | 1.015385 | 1.128866 | 1.041509 | 1.243697 | 1.184615 | 0.974227 | 0.860377 |
| 0.981132 | 1.107692 | 0.912371 | 0.739623 | 1.327731 | 1 | 1.14433 | 1.116981 |
| 1.14717 | 1.092308 | 1.128866 | 1.056604 | 1.142857 | 1.230769 | 1.113402 | 1.056604 |
| 0.996226 | 0.938462 | 1.020619 | 0.981132 | 1.207547 | 1.107692 | 1.14433 | 1.011321 |
| 0.875472 | 1.046154 | 1.121212 | 1.116981 | 1.056604 | 0.876923 | 1.221649 | 1.177358 |
| 0.943089 | 0.984615 | 0.878788 | 1.071698 | 1.177358 | 0.969231 | 1.206186 | 1.086792 |
| 1.056911 | 1.076923 | 0.863636 | 1.14433 | 1.298113 | 1.138462 | 0.742424 | 1.132075 |
| 1.082474 | 1.046154 | 0.848485 | 1.06701 | 1.25283 | 1.2 | 1.19697 | 0.875472 |
| 0.881443 | 1.097938 | 1.121212 | 1.097938 | 1.011321 | 1.138462 | 1.272727 | 0.935849 |
| 1.036082 | 1.113402 | 1.166667 | 1.175258 | 1.298113 | 1.107692 | 1.19697 | 1.221649 |
| 1.045455 | 1.097938 | 1.030303 | 1.075758 | 1.071698 | 1.237113 | 0.772727 | 0.896907 |
| 1.075758 | 1.06701 |  | 0.833333 | 1.298113 | 1.221649 | 0.863636 | 0.909091 |
| 0.80303 | 0.896907 |  | 1.015152 | 1.086792 | 1.190722 | 0.939394 | 0.833333 |
| 1.075758 |  |  | 1.136364 | 1.14717 | 1.376289 | 1.287879 | 1.106061 |
|  |  |  |  | 1.170732 | 1.190722 | 0.863636 | 0.772727 |
|  |  |  |  | 1.235772 | 0.958763 | 1.227273 | 1.166667 |
|  |  |  |  | 1.284553 | 1.314433 | 1.060606 |  |
|  |  |  |  | 1.283505 | 1.036082 | 1.212121 |  |
|  |  |  |  | 1.268041 | 1.268041 |  |  |
|  |  |  |  | 1.051546 | 0.927835 |  |  |
|  |  |  |  | 1.237113 | 1.252577 |  |  |
|  |  |  |  | 1.237113 | 1.175258 |  |  |
|  |  |  |  | 1.082474 |  |  |  |
|  |  |  |  | 1.391753 |  |  |  |
|  |  |  |  | 1.333333 |  |  |  |
|  |  |  |  | 1.333333 |  |  |  |
|  |  |  |  | 1.212121 |  |  |  |

**Table S5:** Raw Data Related to Figure S2b:

| Vehicle | 10µM inhibitor |
| --- | --- |
| 1.768912232 | 0.2980559439 |
| 1.406431678 | 0.5703733404 |
| 1.979891974 | 0.7134650875 |
| 1.94082861 | 0.7286883134 |
| 1.10248142 | 1.03228703 |
| 2.04038843 | 0.6820866272 |
| 1.763995807 | 0.5027542097 |
| 0.4519752608 | 0.8589465744 |
| 0.554655396 | 0.974906994 |
| 0.6014346335 | 0.4181676697 |
| 0.9006407924 | 0.3394453165 |
| 0.4929981929 | 0.8810669726 |
| 1.036008703 | 0.5115177982 |
| 2.258700195 | 0.9094346865 |
| 0.7607533816 | 0.5911148828 |
| 0.6580655943 | 0.4353717814 |
| 0.932866977 | 0.360196572 |
| 0.5751706881 | 0.6759155606 |
| 0.6235023975 | 0.5448133858 |
| 0.6328533306 | 1.659652875 |
| 0.7897882994 | 1.007774796 |
| 0.6006922576 | 0.7858743857 |
| 0.6686981052 | 0.3895679274 |
| 0.8022236996 | 1.033776955 |
| 0.8981248001 | 1.550352538 |
| 0.4481664552 | 1.277899902 |
| 0.6601635715 | 1.415550213 |
| 0.8761503076 | 1.299360521 |
| 0.9032685344 | 2.666303025 |
| 0.6781361046 | 2.42499605 |
| 1.271230475 | 0.9075509585 |
| 0.5710077592 | 0.6923755381 |
| 0.7617371828 | 1.892891999 |
| 1.588056753 | 1.192669706 |
|  | 0.9881308206 |
|  | 1.856820102 |
|  | 2.23577449 |
|  | 0.7922893526 |
|  | 1.274524202 |
|  | 0.8001802708 |

**Table S6:** Raw Data Related to Figure 3f and Figure S3c:

| *nos-Gal4* | *bam-Gal4* | *nos-Gal4ΔVP16; bam-Gal80* | *in Silico nos-Gal4ΔVP16; bam-Gal80 and bam-Gal4* |
| --- | --- | --- | --- |
| 0.833755 | 0.023199 | 3.012224 | 0.023199 |
| 1.205462 | 0.002493 | 1.520095 | 0.002493 |
| 0.067381 | 0.913725 | 0.963087 | 0.913725 |
| 2.185072 | 0.557483 | 0.845611 | 0.557483 |
| 0.596007 | -0.09516 | 0.674583 | -0.09516 |
| 1.240573 | 0.64262 | 1.358367 | 0.64262 |
| 2.138097 | 0.534351 | 2.503381 | 0.534351 |
| 1.509625 | -0.39413 | 1.484098 | -0.39413 |
| 1.578043 | 0.243138 | 0.998465 | 0.243138 |
| -0.70103 | 0.177005 | 0.769789 | 0.177005 |
| 0.720501 | 0.535789 | 2.527987 | 0.535789 |
| 0.441195 | 1.919735 | 0.647042 | 1.919735 |
| 0.055739 | 0.067541 | 2.353069 | 0.067541 |
| -0.92427 | 0.509258 | 2.198496 | 0.509258 |
| 0.628984 | 0.753873 | 1.563693 | 0.753873 |
| 0.870491 | 0.761944 | 1.608681 | 0.761944 |
| 0.460992 | -0.45322 |  | -0.45322 |
| 1.861391 | 0.810568 |  | 0.810568 |
| 1.048597 | -0.50049 |  | -0.50049 |
| 0.120114 | 1.389966 |  | 1.389966 |
| 0.916928 |  |  | 3.012224 |
| 2.28547 |  |  | 1.520095 |
| -0.25156 |  |  | 0.963087 |
| 2.853039 |  |  | 0.845611 |
| 0.400734 |  |  | 0.674583 |
| 2.197903 |  |  | 1.358367 |
| 1.424744 |  |  | 2.503381 |
|  |  |  | 1.484098 |
|  |  |  | 0.998465 |
|  |  |  | 0.769789 |
|  |  |  | 2.527987 |
|  |  |  | 0.647042 |
|  |  |  | 2.353069 |
|  |  |  | 1.563693 |
|  |  |  | 1.608681 |

| *nos-Gal4; polα50<sup>+/-</sup>* | *bam-Gal4; polα50<sup>+/-</sup>* | *nos-Gal4ΔVP16; bam-Gal80; polα50<sup>+/-</sup>* |
| --- | --- | --- |
| 1.71324 | -0.25949 | 1.130421 |
| 1.680616 | 0.614043 | 1.889422 |
| 1.085999 | 1.229249 | 1.55281 |
| 1.620811 | 0.831694 | 0.938152 |
| 1.004243 | 1.557561 | 1.519162 |
| 1.134904 | 1.30331 | 0.997146 |
| 0.538452 | 1.387124 | 1.289368 |
| 1.377369 | 0.413369 | 0.700357 |
| 0.209853 | 2.71423 | 0.66006 |
| 1.277605 | 0.927196 | 1.754169 |
| 0.165427 | 0.02529 | 2.32908 |
| 2.37251 | 0.590838 | 0.938002 |
| 1.188624 | 1.190953 | 2.784909 |
| 1.429034 | 1.138515 | 1.103144 |
| 2.772322 | 1.336803 | 2.021303 |
| 3.005586 | 2.862607 | 2.376666 |
| 2.465316 | 0.426856 | 1.520804 |
| 1.284234 | 0.29905 | 0.642919 |
| 1.706529 | 1.812977 | 1.529886 |
| 1.52534 | 3.308225 | 1.460099 |
| 1.517268 | -0.05976 | 1.396838 |
| 1.403463 | 1.058061 | 1.719726 |
| 1.667895 |  |  |

**Table S7:**
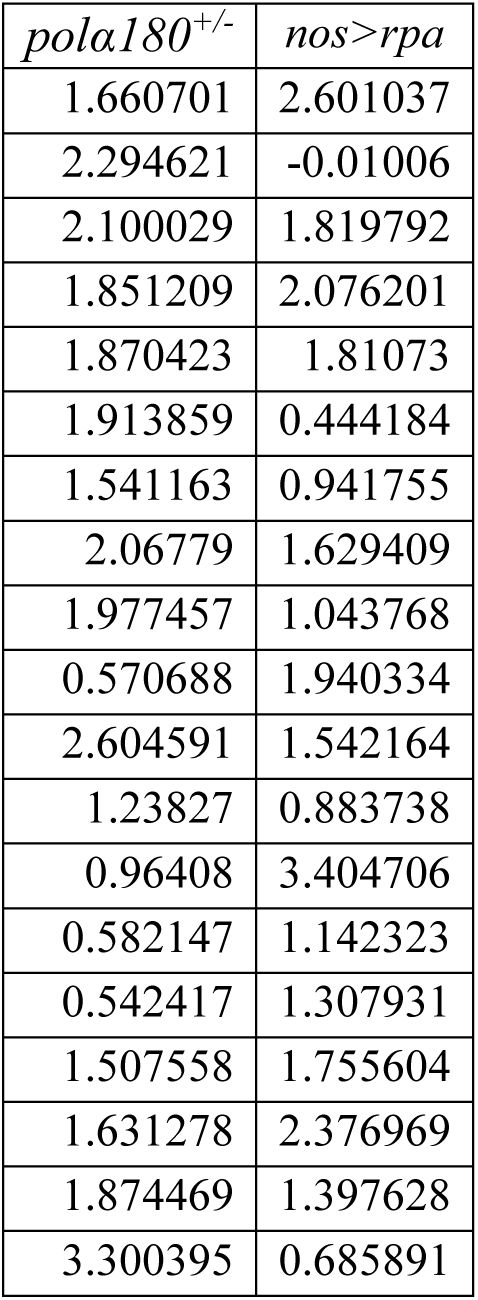
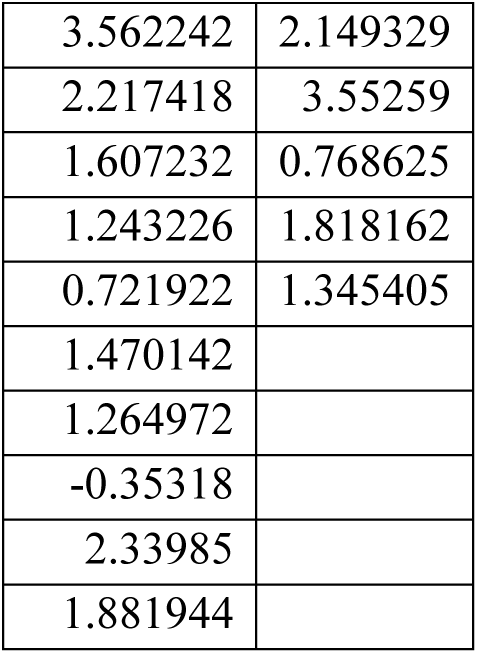
Raw Data Related to Figure S3d:

**Table S8:** Raw Data Related to Figure 4e: Compaction index using log_2_ scale in control and *pola50^+/-^* GSCs and 8-cell SGs.

| GSC | SG | *pola50*<sup>+/-</sup> GSC | *pola50*<sup>+/-</sup> SG |
| --- | --- | --- | --- |
| 1.051 | 0.581 | 1.868 | 1.409 |
| 1.948 | 0.210 | 0.539 | 1.825 |
| 1.070 | 0.658 | 1.358 | 1.233 |
| 0.966 | 0.554 | 0.814 | 1.410 |
| 1.588 | 0.507 | 0.787 | 1.692 |
| 0.959 | 0.565 | 0.607 | 0.842 |
| 1.491 | 0.215 | 1.330 | 2.127 |
| 1.166 | 0.129 | 1.992 | 2.262 |
| 1.859 | 0.515 | 0.790 | 1.478 |
| 0.679 | 0.369 | 2.284 | 1.312 |
| 1.376 | 0.370 | 0.792 | 0.747 |
| 2.800 | 0.201 | 0.975 |  |
| 0.676 |  | 1.616 |  |
| 0.831 |  |  |  |
| 0.557 |  |  |  |

**Table S9:** Related to Figure S4a:

| Symmetric (EdU/EdU fiber) | Weak Asymmetry (EdU/BrdU less than 2-fold asymmetry) | Strong Asymmetry (EdU/BrdU with greater than 2-fold asymmetry) |
| --- | --- | --- |
| 31 | 12 | 9 |

**Table S10:** Raw Data Related to Figure 5c and Figure S4b:

| Log<sub>2</sub>(BrdU) | Log<sub>2</sub>(EdU) |
| --- | --- |
| 0.949727 | -0.78622 |
| 3.979822 | -2.37851 |
| 2.947978 | -3.12338 |
| 0.975338 | -0.26303 |
| 0.494109 | -0.52319 |
| 1.794681 | -0.69707 |
| 0.313499 | -0.90752 |
| 0.264416 | 0.147226 |
| 0.285157 | -0.40526 |
| 2.560361 | -1.06054 |
| 0.377185 | -0.94404 |
| 2.900464 | -1.48543 |
| 0.385654 | -0.24246 |
| 1.201369 | -1.25831 |
| 1.076044 | -0.61857 |
| 1.662965 | -1.5025 |
| 1.730393 | 0.21864 |
| 0.171511 | -0.98296 |
| 0.587198 | -0.9783 |
| 0.373352 | -0.69574 |
| 0.830075 | -1.11346 |

**Table S11:** Log_2_ Raw Data Related to Figure 5d and Figure S4f*:

| *nos-Gal4* | *nos-Gal4;*<br>*polα50<sup>+/-</sup>* | *bam-Gal4* | *bam-Gal4;*<br>*polα50<sup>+/-</sup>* | *nos-*<br>*Gal4ΔVP16;*<br>*bam-Gal80* | *nos-Gal4 ΔVP16;*<br>*bam-Gal80;*<br>*polα50<sup>+/-</sup>* |
| --- | --- | --- | --- | --- | --- |
| -1.64051 | 1.706293 | 0.539826 | 0.456845 | -0.73765 | -1.36482 |
| 2.612107 | -3.25913 | 0.140349 | -1.57106 | -0.95258 | 0.75138 |
| 0.451143 | 0.354129 | 0.035564 | -1.05006 | -1.89181 | -0.03933 |
| -1.74333 | 1.783676 | -1.15265 | -0.75268 | -1.1849 | 0.70204 |
| 0.546183 | -1.6443 | -0.31179 | 0.033124 | 2.108614 | -1.28129 |
| -2.88245 | -0.65438 | 0.959769 | 0.805337 | -1.90253 | -1.04057 |
| -3.72782 | -2.72158 | -0.0864 | -2.00483 | -1.708 | -0.73152 |
| -1.44057 | -3.32288 | -0.47096 | -2.517 | -1.46108 | -3.05472 |
| -3.62121 | -0.39184 | 0.849896 | -0.83188 | -4.2996 | -2.8483 |
| -2.73246 | -0.87832 | -0.03163 | -0.31128 | -1.11643 | -2.33536 |
| 0.096365 | -0.95884 | 0.505262 | -1.53716 | 0.266644 | -3.35477 |
| 0.701662 | -1.14961 | -0.1108 | -0.42537 | -0.20308 | -1.95236 |
| 2.49744 | 0.442487 | 0.371136 | 0.274047 | -1.95157 | -1.36273 |
| -0.17693 | -2.98338 | 0.388789 | -1.43808 | 1.353261 | -1.56238 |
| 0.415777 | -0.97708 | 0.515224 | -0.99389 | -0.96038 | 0.908932 |
| 0.63151 | -1.30605 | -0.28391 | -0.84444 | -1.6619 | -1.53928 |
| -0.7567 | 1.10949 | 0.62232 | -0.5031 | 2.020612 | -2.43426 |
| -0.99991 | 1.181099 | -1.15803 | -1.97743 | -1.48487 | -3.2655 |
| -1.29837 | 1.226325 | -0.03333 | -1.83919 |  | 0.490629 |
| 0.530978 | -1.62792 | -0.69519 | -1.66125 |  | -1.31617 |
| -1.07129 | -2.34696 | -1.28691 | -0.6374 |  | -0.80856 |
| -0.36712 | -1.43265 |  | -0.91854 |  |  |
| 0.237249 | -1.42746 |  | 1.550994 |  |  |
| -1.4912 | -1.52134 |  |  |  |  |
| 1.076845 | -3.97101 |  |  |  |  |
| -0.37702 | -2.85406 |  |  |  |  |
| 0.469742 |  |  |  |  |  |

(\*Data are provided in absolute values in Fig.5d and log<sub>2</sub> values in Fig.S4f)

**Table S12:** Raw Data Related to Figure 5g:

| Polα | Polε |
| --- | --- |
| 0.581 | 0.531 |
| 4.038 | 0.6 |
| 7.409 | 0.497 |
| 1.922 | 0.368 |
| 2.366 | 1.269 |
| 2.908 | 0.698 |
| 3.641 | 0.822 |
| 0.983 | 0.727 |
| 1.577 | 0.776 |
| 3.312 | 0.316 |
| 3.657 | 0.908 |
| 1.293 | 0.309 |
| 2.333 | 0.223 |
| 8.184 | 0.161 |
|  | 0.147 |

**Table S13:** Related to Figure S4c:

| Strong asymmetry toward the leading strand ($\geq 2$-fold) | Weak asymmetry toward the leading stand ($< 2$-fold) | Weak asymmetry toward the lagging stand ($< 2$-fold) | Strong asymmetry toward the lagging strand ($\geq 2$-fold) |
| --- | --- | --- | --- |
| 3 | 9 | 4 | 11 |

**Table S14:** Related to Figure S4d:

| PCNA | EdU | H3K27me3 |
| --- | --- | --- |
| -1.05482 | -1.64051 | 0.833755 |
| -1.21579 | 2.612107 | 1.205462 |
| -0.3613 | 0.451143 | 0.067381 |
| -0.85911 | -1.74333 | 2.185072 |
| -0.7166 | 0.546183 | 0.596007 |
| -5.24247 | -2.88245 | 1.240573 |
| -1.49709 | -3.72782 | 2.138097 |
| -0.56181 | -1.44057 | 1.509625 |
| -2.67029 | -3.62121 | 1.578043 |
| -3.0681 | -2.73246 | -0.70103 |
| -0.49276 | 0.096365 | 0.720501 |
| -1.1499 | 0.701662 | 0.441195 |
| -0.82622 | 2.49744 | 0.055739 |
| -0.12406 | -0.17693 | -0.92427 |
| -1.09902 | 0.415777 | 0.628984 |
| -0.99608 | 0.63151 | 0.870491 |
| -1.45258 | -0.7567 | 0.460992 |
| -2.06082 | -0.99991 | 1.861391 |
| -0.93227 | -1.29837 | 1.048597 |
| -1.53805 | 0.530978 | 0.120114 |
| -0.83251 | -1.07129 | 0.916928 |
| -3.9494 | -0.36712 | 2.28547 |
| -3.90744 | 0.237249 | -0.25156 |
| -1.45679 | -1.4912 | 2.853039 |
| -0.33855 | 1.076845 | 0.400734 |
| -1.61222 | -0.37702 | 2.197903 |
| -1.04414 | 0.469742 | 1.424744 |

## References

1. C. D. Allis, T. Jenuwein, The molecular hallmarks of epigenetic control. Nat Rev Genet 17, 487–500 (2016).

2. A. D. Goldberg, C. D. Allis, E. Bernstein, Epigenetics: a landscape takes shape. Cell 128, 635–638 (2007).

3. R. Bonasio, S. Tu, D. Reinberg, Molecular signals of epigenetic states. Science 330, 612–616 (2010).

4. T. M. Escobar, A. Loyola, D. Reinberg, Parental nucleosome segregation and the inheritance of cellular identity. Nat Rev Genet 22, 379–392 (2021).

5. J. A. Urban, R. Ranjan, X. Chen, Asymmetric Histone Inheritance: Establishment, Recognition, and Execution. Annu Rev Genet, (2022).

6. S. I. S. Grewal, The molecular basis of heterochromatin assembly and epigenetic inheritance. Mol Cell 83, 1767–1785 (2023).

7. A. E. Vouzas, D. M. Gilbert, Replication timing and transcriptional control: beyond cause and effect - part IV. Curr Opin Genet Dev 79, 102031 (2023).

8. B. Sunchu, C. Cabernard, Principles and mechanisms of asymmetric cell division. Development 147, (2020).

9. Z. G. Venkei, Y. M. Yamashita, Emerging mechanisms of asymmetric stem cell division. J Cell Biol 217, 3785–3795 (2018).

10. E. H. Zion, C. Chandrasekhara, X. Chen, Asymmetric inheritance of epigenetic states in asymmetrically dividing stem cells. Curr Opin Cell Biol 67, 27–36 (2020).

11. C. Blanpain, E. Fuchs, Stem cell plasticity. Plasticity of epithelial stem cells in tissue regeneration. Science 344, 1242281 (2014).

12. J. A. Knoblich, Asymmetric cell division: recent developments and their implications for tumour biology. Nat Rev Mol Cell Biol 11, 849–860 (2010).

13. J. Ferrand et al., Mitotic chromatin marking governs asymmetric segregation of DNA damage. bioRxiv, (2023).

14. V. Tran, C. Lim, J. Xie, X. Chen, Asymmetric division of Drosophila male germline stem cell shows asymmetric histone distribution. Science 338, 679–682 (2012).

15. M. Wooten et al., Asymmetric histone inheritance via strand-specific incorporation and biased replication fork movement. Nat Struct Mol Biol 26, 732–743 (2019).

16. E. H. Zion et al., Old and newly synthesized histones are asymmetrically distributed in Drosophila intestinal stem cell divisions. EMBO Rep 24, e56404 (2023).

17. M. Wooten et al., Superresolution imaging of chromatin fibers to visualize epigenetic information on replicative DNA. Nat Protoc 15, 1188–1208 (2020).

18. R. Ranjan, J. Snedeker, X. Chen, Asymmetric Centromeres Differentially Coordinate with Mitotic Machinery to Ensure Biased Sister Chromatid Segregation in Germline Stem Cells. Cell Stem Cell 25, 666–681 e665 (2019).

19. R. Ranjan et al., Differential condensation of sister chromatids acts with Cdc6 to ensure asynchronous S-phase entry in Drosophila male germline stem cell lineage. Dev Cell 57, 1102–1118 e1107 (2022).

20. M. Antel et al., Interchromosomal interaction of homologous Stat92E alleles regulates transcriptional switch during stem-cell differentiation. Nat Commun 13, 3981 (2022).

21. C. Yu et al., A mechanism for preventing asymmetric histone segregation onto replicating DNA strands. Science 361, 1386–1389 (2018).

22. H. Gan et al., The Mcm2-Ctf4-Polalpha Axis Facilitates Parental Histone H3-H4 Transfer to Lagging Strands. Mol Cell 72, 140–151 e143 (2018).

23. Z. Shan et al., The patterns and participants of parental histone recycling during DNA replication in Saccharomyces cerevisiae. Sci China Life Sci, (2023).

24. G. Schlissel, J. Rine, The nucleosome core particle remembers its position through DNA replication and RNA transcription. Proc Natl Acad Sci U S A 116, 20605–20611 (2019).

25. N. Li et al., Parental histone transfer caught at the replication fork. Nature 627, 890–897 (2024).

26. N. Petryk et al., MCM2 promotes symmetric inheritance of modified histones during DNA replication. Science 361, 1389–1392 (2018).

27. T. M. Escobar et al., Active and Repressed Chromatin Domains Exhibit Distinct Nucleosome Segregation during DNA Replication. Cell 179, 953–963 e911 (2019).

28. Z. Li et al., DNA polymerase alpha interacts with H3-H4 and facilitates the transfer of parental histones to lagging strands. Sci Adv 6, eabb5820 (2020).

29. A. Wenger et al., Symmetric inheritance of parental histones governs epigenome maintenance and embryonic stem cell identity. Nat Genet 55, 1567–1578 (2023).

30. V. Flury et al., Recycling of modified H2A-H2B provides short-term memory of chromatin states. Cell 186, 1050–1065 e1019 (2023).

31. N. Reveron-Gomez et al., Accurate Recycling of Parental Histones Reproduces the Histone Modification Landscape during DNA Replication. Mol Cell 72, 239–249 e235 (2018).

32. M. Xu et al., Partitioning of histone H3-H4 tetramers during DNA replication-dependent chromatin assembly. Science 328, 94–98 (2010).

33. W. Du et al., Mechanisms of chromatin-based epigenetic inheritance. Sci China Life Sci 65, 2162–2190 (2022).

34. A. Serra-Cardona, Z. Zhang, Replication-Coupled Nucleosome Assembly in the Passage of Epigenetic Information and Cell Identity. Trends Biochem Sci 43, 136–148 (2018).

35. K. R. Stewart-Morgan, N. Petryk, A. Groth, Chromatin replication and epigenetic cell memory. Nat Cell Biol 22, 361–371 (2020).

36. W. Zhang, J. Feng, Q. Li, The replisome guides nucleosome assembly during DNA replication. Cell Biosci 10, 37 (2020).

37. M. D. White et al., Long-Lived Binding of Sox2 to DNA Predicts Cell Fate in the Four-Cell Mouse Embryo. Cell 165, 75–87 (2016).

38. D. a. R. Wen, Z., Activation of the Maternal Genome Through Asymmetric Distribution of Oocyte-Genome-Associated Histone H3.3. bioRxiv, (2023).

39. M. Foltman et al., Eukaryotic replisome components cooperate to process histones during chromosome replication. Cell Rep 3, 892–904 (2013).

40. T. Iida, H. Araki, Noncompetitive counteractions of DNA polymerase epsilon and ISW2/yCHRAC for epigenetic inheritance of telomere position effect in Saccharomyces cerevisiae. Mol Cell Biol 24, 217–227 (2004).

41. D. S. Saxton, J. Rine, Epigenetic memory independent of symmetric histone inheritance. Elife 8, (2019).

42. P. G. Mitsis, S. C. Kowalczykowski, I. R. Lehman, A single-stranded DNA binding protein from Drosophila melanogaster: characterization of the heterotrimeric protein and its interaction with single-stranded DNA. Biochemistry 32, 5257–5266 (1993).

43. R. F. Marton, P. Thommes, S. Cotterill, Purification and characterisation of dRP-A: a single-stranded DNA binding protein from Drosophila melanogaster. FEBS Lett 342, 139–144 (1994).

44. S. A. Blythe, E. F. Wieschaus, Zygotic genome activation triggers the DNA replication checkpoint at the midblastula transition. Cell 160, 1169–1181 (2015).

45. J. Chen, S. Le, A. Basu, W. J. Chazin, J. Yan, Mechanochemical regulations of RPA’s binding to ssDNA. Scientific Reports 5, 9296 (2015).

46. M. L. Jones, V. Aria, Y. Baris, J. T. P. Yeeles, How Pol alpha-primase is targeted to replisomes to prime eukaryotic DNA replication. Mol Cell 83, 2911–2924 e2916 (2023).

47. M. R. G. Taylor, J. T. P. Yeeles, The Initial Response of a Eukaryotic Replisome to DNA Damage. Mol Cell 70, 1067–1080 e1012 (2018).

48. K. L. Collins, T. J. Kelly, Effects of T antigen and replication protein A on the initiation of DNA synthesis by DNA polymerase alpha-primase. Mol Cell Biol 11, 2108–2115 (1991).

49. H. Huang et al., Structure of a DNA polymerase alpha-primase domain that docks on the SV40 helicase and activates the viral primosome. J Biol Chem 285, 17112–17122 (2010).

50. K. Weisshart et al., Protein-protein interactions of the primase subunits p58 and p48 with simian virus 40 T antigen are required for efficient primer synthesis in a cell-free system. J Biol Chem 275, 17328–17337 (2000).

51. P. Gadre, N. Nitsure, D. Mazumdar, S. Gupta, K. Ray, The rates of stem cell division determine the cell cycle lengths of its lineage. iScience 24, 103232 (2021).

52. T. D. Carroll, I. P. Newton, Y. Chen, J. J. Blow, I. Nathke, Lgr5(+) intestinal stem cells reside in an unlicensed G1 phase. J Cell Biol 217, 1667–1685 (2018).

53. R. Ranjan, X. Chen, Quantitative imaging of chromatin inheritance using a dual-color histone in Drosophila germinal stem cells. STAR Protoc 3, 101811 (2022).

54. M. Sivaguru et al., Comparative performance of airyscan and structured illumination superresolution microscopy in the study of the surface texture and 3D shape of pollen. Microsc Res Tech 81, 101–114 (2018).

55. E. W. Kahney et al., Characterization of histone inheritance patterns in the Drosophila female germline. EMBO Rep, e51530 (2021).

56. C. Chandrasekhara et al., A single N-terminal amino acid determines the distinct roles of histones H3 and H3.3 in the Drosophila male germline stem cell lineage. PLoS Biol 21, e3002098 (2023).

57. R. Cincinelli et al., Novel adamantyl retinoid-related molecules with POLA1 inhibitory activity. Bioorg Chem 104, 104253 (2020).

58. M. Inaba, M. Buszczak, Y. M. Yamashita, Nanotubes mediate niche-stem-cell signalling in the Drosophila testis. Nature 523, 329–332 (2015).

59. N. R. Matias, J. Mathieu, J. R. Huynh, Abscission is regulated by the ESCRT-III protein shrub in Drosophila germline stem cells. PLoS Genet 11, e1004653 (2015).

60. M. Van Doren, A. L. Williamson, R. Lehmann, Regulation of zygotic gene expression in Drosophila primordial germ cells. Curr Biol 8, 243–246 (1998).

61. S. H. Eun, Z. Shi, K. Cui, K. Zhao, X. Chen, A non-cell autonomous role of E(z) to prevent germ cells from turning on a somatic cell marker. Science 343, 1513–1516 (2014).

62. J. Cheng et al., Centrosome misorientation reduces stem cell division during ageing. Nature 456, 599–604 (2008).

63. D. Chen, D. M. McKearin, A discrete transcriptional silencer in the bam gene determines asymmetric division of the Drosophila germline stem cell. Development 130, 1159–1170 (2003).

64. C. Alabert et al., Two distinct modes for propagation of histone PTMs across the cell cycle. Genes Dev 29, 585–590 (2015).

65. S. Lin, Z. F. Yuan, Y. Han, D. M. Marchione, B. A. Garcia, Preferential Phosphorylation on Old Histones during Early Mitosis in Human Cells. J Biol Chem 291, 15342–15357 (2016).

66. C. Yu et al., Strand-specific analysis shows protein binding at replication forks and PCNA unloading from lagging strands when forks stall. Mol Cell 56, 551–563 (2014).

67. S. Petruk et al., TrxG and PcG proteins but not methylated histones remain associated with DNA through replication. Cell 150, 922–933 (2012).

68. T. K. Fenstermaker, S. Petruk, S. K. Kovermann, H. W. Brock, A. Mazo, RNA polymerase II associates with active genes during DNA replication. Nature 620, 426–433 (2023).

69. J. E. Graham, K. J. Marians, S. C. Kowalczykowski, Independent and Stochastic Action of DNA Polymerases in the Replisome. Cell 169, 1201–1213 e1217 (2017).

70. A. Ercilla et al., Physiological Tolerance to ssDNA Enables Strand Uncoupling during DNA Replication. Cell Rep 30, 2416–2429 e2417 (2020).

71. R. Ziane, A. Camasses, M. Radman-Livaja, The asymmetric distribution of RNA polymerase II and nucleosomes on replicated daughter genomes is caused by differences in replication timing between the lagging and the leading strand. Genome Res 32, 337–356 (2022).

72. M. L. Bochman, A. Schwacha, The Mcm complex: unwinding the mechanism of a replicative helicase. Microbiol Mol Biol Rev 73, 652–683 (2009).

73. B. K. Tye, MCM proteins in DNA replication. Annu Rev Biochem 68, 649–686 (1999).

74. L. D. Brennan, R. A. Forties, S. S. Patel, M. D. Wang, DNA looping mediates nucleosome transfer. Nat Commun 7, 13337 (2016).

75. Y. M. Yamashita, A. P. Mahowald, J. R. Perlin, M. T. Fuller, Asymmetric inheritance of mother versus daughter centrosome in stem cell division. Science 315, 518–521 (2007).

76. B. Arezi, R. D. Kuchta, Eukaryotic DNA primase. Trends Biochem Sci 25, 572–576 (2000).

77. E. Johansson, S. A. Macneill, The eukaryotic replicative DNA polymerases take shape. Trends Biochem Sci 35, 339–347 (2010).

78. Z. X. Zhou, S. A. Lujan, A. B. Burkholder, M. A. Garbacz, T. A. Kunkel, Roles for DNA polymerase delta in initiating and terminating leading strand DNA replication. Nat Commun 10, 3992 (2019).

79. R. E. Johnson, R. Klassen, L. Prakash, S. Prakash, A Major Role of DNA Polymerase delta in Replication of Both the Leading and Lagging DNA Strands. Mol Cell 59, 163–175 (2015).

80. S. Liu et al., RPA binds histone H3-H4 and functions in DNA replication-coupled nucleosome assembly. Science 355, 415–420 (2017).

81. R. Cincinelli et al., A novel atypical retinoid endowed with proapoptotic and antitumor activity. J Med Chem 46, 909–912 (2003).

82. R. R. Nasr et al., ST1926, an orally active synthetic retinoid, induces apoptosis in chronic myeloid leukemia cells and prolongs survival in a murine model. Int J Cancer 137, 698–709 (2015).

83. M. R. H. Zwinderman et al., Deposition Bias of Chromatin Proteins Inverts under DNA Replication Stress Conditions. ACS Chem Biol 16, 2193–2201 (2021).

84. M. M. Seidman, A. J. Levine, H. Weintraub, The asymmetric segregation of parental nucleosomes during chrosome replication. Cell 18, 439–449 (1979).

85. Z. Li et al., Asymmetric distribution of parental H3K9me3 in S phase silences L1 elements. Nature, (2023).

86. T. Nakatani et al., DNA replication fork speed underlies cell fate changes and promotes reprogramming. Nat Genet 54, 318–327 (2022).

87. T. Nakatani et al., Emergence of replication timing during early mammalian development. Nature 625, 401–409 (2024).

88. K. N. Klein et al., Replication timing maintains the global epigenetic state in human cells. Science 372, 371–378 (2021).

89. J. Sima et al., Identifying cis Elements for Spatiotemporal Control of Mammalian DNA Replication. Cell 176, 816–830 e818 (2019).

90. D. Muhlen, X. Li, O. Dovgusha, H. Jackle, U. Gunesdogan, Recycling of parental histones preserves the epigenetic landscape during embryonic development. Sci Adv 9, eadd6440 (2023).

91. W. Hennig, A. Weyrich, Histone modifications in the male germ line of Drosophila. BMC Dev Biol 13, 7 (2013).

92. J. Xie et al., Histone H3 Threonine Phosphorylation Regulates Asymmetric Histone Inheritance in the Drosophila Male Germline. Cell 163, 920–933 (2015).

## Supplemental References

1. M. Van Doren, A. L. Williamson, R. Lehmann, Regulation of zygotic gene expression in Drosophila primordial germ cells. Curr Biol 8, 243–246 (1998).

2. M. Inaba, M. Buszczak, Y. M. Yamashita, Nanotubes mediate niche-stem-cell signalling in the Drosophila testis. Nature 523, 329–332 (2015).

3. D. Chen, D. M. McKearin, A discrete transcriptional silencer in the bam gene determines asymmetric division of the Drosophila germline stem cell. Development 130, 1159–1170 (2003).

4. M. Wooten et al., Asymmetric histone inheritance via strand-specific incorporation and biased replication fork movement. Nat Struct Mol Biol 26, 732–743 (2019).

5. S. A. Blythe, E. F. Wieschaus, Zygotic genome activation triggers the DNA replication checkpoint at the midblastula transition. Cell 160, 1169–1181 (2015).

6. V. Tran, C. Lim, J. Xie, X. Chen, Asymmetric division of Drosophila male germline stem cell shows asymmetric histone distribution. Science 338, 679–682 (2012).

7. T. D. Carroll, I. P. Newton, Y. Chen, J. J. Blow, I. Nathke, Lgr5(+) intestinal stem cells reside in an unlicensed G1 phase. J Cell Biol 217, 1667–1685 (2018).

8. R. Ranjan et al., Differential condensation of sister chromatids acts with Cdc6 to ensure asynchronous S-phase entry in Drosophila male germline stem cell lineage. Dev Cell 57, 1102–1118 e1107 (2022).

9. R. Ranjan, J. Snedeker, X. Chen, Asymmetric Centromeres Differentially Coordinate with Mitotic Machinery to Ensure Biased Sister Chromatid Segregation in Germline Stem Cells. Cell Stem Cell 25, 666–681 e665 (2019).

10. J. Cheng et al., Centrosome misorientation reduces stem cell division during ageing. Nature 456, 599–604 (2008).

11. Y. M. Yamashita, D. L. Jones, M. T. Fuller, Orientation of asymmetric stem cell division by the APC tumor suppressor and centrosome. Science 301, 1547–1550 (2003).

12. Y. M. Yamashita, A. P. Mahowald, J. R. Perlin, M. T. Fuller, Asymmetric inheritance of mother versus daughter centrosome in stem cell division. Science 315, 518–521 (2007).

13. S. Yadlapalli, J. Cheng, Y. M. Yamashita, Drosophila male germline stem cells do not asymmetrically segregate chromosome strands. J Cell Sci 124, 933–939 (2011).

14. X. R. Sheng, E. Matunis, Live imaging of the Drosophila spermatogonial stem cell niche reveals novel mechanisms regulating germline stem cell output. Development 138, 3367–3376 (2011).

15. M. Wooten et al., Superresolution imaging of chromatin fibers to visualize epigenetic information on replicative DNA. Nat Protoc 15, 1188–1208 (2020).

16. R. Cincinelli et al., Novel adamantyl retinoid-related molecules with POLA1 inhibitory activity. Bioorg Chem 104, 104253 (2020).

